# Maternal genome dominance in early plant embryogenesis

**DOI:** 10.1101/2020.01.14.905992

**Authors:** Jaime Alaniz-Fabián, Daoquan Xiang, Gerardo Del Toro-De León, Axel Orozco-Nieto, Peng Gao, Andrew Sharpe, Leon V. Kochian, Gopalan Selvaraj, Nathan Springer, Cei Abreu-Goodger, Raju Datla, C. Stewart Gillmor

## Abstract

Previous studies have alternately supported and discounted the hypothesis that the maternal genome plays a predominant role in early embryogenesis in plants. We used 24 *embryo defective (emb)* mutants of *Arabidopsis thaliana* to test for maternal and paternal effects in early embryogenesis. 5 *emb* mutants had equal maternal and paternal effects, 5 showed maternal effects and weak paternal effects, and the remaining 14 *emb* mutants conditioned only maternal effects, demonstrating a more important role for the maternal allele for most of these *EMB* genes. To assess genome-wide maternal and paternal contributions to early embryos, we produced allele-specific transcriptomes from zygote to mature stage embryos derived from reciprocal crosses of Columbia-0 and Tsu-1, a hybrid combination we show to be a faithful proxy for isogenic Columbia-0. Parent-of-origin analysis of these transcriptomes revealed a reciprocal maternal bias in thousands of genes from the zygote to octant stage. This bias greatly diminished by the globular stage, and was absent at later stages. Comparison with egg cell transcriptomes revealed no correlation between transcript levels in the egg and maternal bias in pre-globular embryos, suggesting that the maternal bias observed in early embryos is due to preferential zygotic transcription of maternal alleles. Taken together, the functional and transcriptome data presented here support a predominant role for the maternal genome in early Arabidopsis embryogenesis.

**Significance:** In both animals and plants, the zygote is produced by the union of the egg and sperm cells. In animals, it is well accepted that mRNAs and proteins from the egg direct the first steps of embryogenesis. Here we present genetic and genomic experiments that support a predominant role for the maternal genome in early embryogenesis of plants, as well. In contrast to animals, our data suggest that this maternal influence is primarily derived not from inheritance of egg transcripts, but from preferential transcription of maternal alleles in the zygote and early embryo. This transient maternal zygotic bias may reflect an ancestral condition to diminish paternal influence on early embryogenesis in outcrossing plants.

## Introduction

In animal embryogenesis, mRNAs and proteins inherited from the egg regulate early development until maternal products are cleared and the zygotic genome is activated, a process called the maternal to zygotic transition (Tadros & Lipshitz, 2009). In plants, most research on the maternal to zygotic transition has focused on activation of the zygotic genome. The main questions currently debated are the timing of initiation of large scale zygotic transcription, and whether this transcription initially has a maternal bias (reviewed in Del Toro-De León et al., 2016).

Initial experiments to study the onset of transcription after fertilization in wheat, maize and tobacco found novel transcripts in cDNA libraries of late zygotes compared to the egg cell (Sauter et al.,1998; Okamoto et al., 2005; Sprunck et al., 2005; Zhao et al., 2011). Recent RNA sequencing experiments comparing gametes with isogenic embryos in maize, rice and Arabidopsis have provided more evidence for transcription of a portion of the zygotic genome within hours after fertilization. In maize zygotes at 12 hours after pollination (hap) (corresponding to 4 hours after fertilization (haf) and to the G1/S phase of the cell cycle), 3605 genes were found to be upregulated compared to the egg and sperm (Chen et al., 2017). In rice, where fertilization occurs about 30 minutes after pollination, comparison of zygote and egg cell transcriptomes found 499 genes upregulated at least two-fold in zygotes at 2.5 hap (corresponding to karyogamy), 1981 genes at 5 hap (nucleolar fusion), and 2485 genes at 9 hap (G2 phase) (Anderson et al., 2017). Arabidopsis zygotes at 14 hap (approximately 6 haf) showed 2625 genes upregulated at least two-fold compared to egg cells, while at 24 hap 2951 genes were upregulated compared to egg cells (Zhao et al., 2019). In summary, these experiments show transcriptional activity for about 10% of the genome in isogenic zygotes of various plant species.

Initial experiments on maternal and paternal contributions to early isogenic embryos relied on reporter genes and functional analysis of early-acting *embryo defective (emb)* mutations. Assays of maternally and paternally-inherited reporter lines in Arabidopsis found almost 30 genes expressed primarily from the maternal allele (Vielle-Calzada et al., 2000; Autran et al., 2011; Del Toro-De León et al., 2014), as well as numerous genes with more equal maternal and paternal expression or function (Weijers et al., 2001; Lukowitz et al., 2004; Xu et al., 2005; Andreuzza et al., 2010; Aw et al., 2010; Ueda et al., 2011; Guo et al., 2016; Yang et al., 2017). In Arabidopsis, the functions of hundreds of *EMBRYO DEFECTIVE (EMB)* genes are required in early embryogenesis (Meinke, 2019). Crosses of 49 *emb* mothers to isogenic wild type plants showed that paternal alleles for 9 *EMB* genes immediately complemented lack of maternal function, while paternal alleles for 40 *EMB* genes did not fully complement until 3-5 dap (Del Toro-De León et al., 2014). Consistent with the RNA sequencing experiments mentioned above, these functional genetic experiments in isogenic embryos show that zygotic activation varies between genes, with much of the paternal genome showing less activity than the maternal genome in the first 2-3 dap.

Maternal and paternal transcripts in embryos can also be quantified using single nucleotide polymorphisms generated by crossing polymorphic ecotypes. Initial allele-specific expression (ASE) RNA sequencing (RNAseq) experiments in Arabidopsis found evidence for a transient maternal bias in early Landsberg *erecta* (L*er*) x Columbia-0 (Col) embryos (Autran et al., 2011), and for early and equal maternal and paternal contributions in Col x Cape Verde Islands (Cvi) embryos (Nodine and Bartel, 2012). Recent ASE RNAseq experiments of hybrid zygotes have also included data for gametes, allowing for comparison of the egg, sperm and zygote transcriptomes. In rice, more than 97% of 14,049 genes had a strong maternal bias at both 2.5 and 9 hap, with maternal transcripts for the vast majority of genes found in both the egg and zygote (Anderson et al., 2017). In Arabidopsis, Col x L*er* hybrid zygotes had a moderate maternal bias at 14 hap (9.9% of 12,746 genes showed maternally biased transcripts), but no maternal bias by 24 hap (Zhao et al., 2019). 60% of upregulated genes at 24 hap showed biallelic expression, suggesting that maternal and paternal alleles of many genes are equally transcribed in late Col x L*er* zygotes (Zhao et al., 2019), similar to Col x Cvi embryos (Nodine and Bartel, 2012). In summary, ASE experiments in hybrid rice zygotes (Anderson et al., 2017) and Arabidopsis L*er* x Col embryos (Autran et al., 2011) argue for primarily maternal regulation of early embryogenesis, while the results of subsequent Col x L*er* (Zhao et al., 2019) and Col x Cvi (Nodine and Bartel, 2012) transcriptomes in Arabidopsis favor both maternal and paternal (i.e. zygotic) regulation of early embryogenesis. Given the differing conclusions of these studies, the extent of maternal and zygotic regulation of early embryogenesis requires further clarification.

To test whether different combinations of genetic backgrounds might explain the divergent results of hybrid transcriptomes, we previously assayed maternal and paternal effects for 11 *emb* mutants in isogenic and hybrid crosses. Maternal effects, but no paternal effects, were observed for all 11 *emb* mutants. In the embryos produced by Col *emb/*+ mothers, the paternal *EMB* allele was activated significantly earlier when it was provided by L*er* or Cvi pollen compared to Wassilewskija, Tsu-1 (Tsu) or Col pollen (Del Toro-Del León et al., 2014). Here, we tested maternal and paternal effects for 24 additional *emb* mutants. These new experiments provided more evidence that maternal effects are more common than paternal effects in early embryogenesis, and that paternal allele activation is similar between the Col x Tsu hybrid and isogenic Col. We used a hand dissection protocol for isolation of zygote-1 cell to mature stage embryos from bi-directional crosses of Col x Tsu, and analyzed maternal and paternal contributions to the transcriptome. Our results demonstrate a maternal transcript bias for thousands of genes in preglobular Col x Tsu and Tsu x Col embryos, consistent with the predominant maternal effects observed in functional genetic studies with *emb* mutants. Interestingly, maternal transcript bias of genes in zygote to octant stage embryos was not correlated with the relative expression of the corresponding gene in the egg cell. Analysis of intronic reads for several hundred genes at the zygote and octant stages demonstrated that the majority of transcripts come from the maternal allele. Taken together, these results support a predominant role for the maternal genome in early embryogenesis, and suggest that this maternal dominance is due more to preferential transcription of maternal alleles in the zygote than to inheritance of maternal mRNAs from the egg cell.

## Results and Discussion

### Maternal effects predominate over paternal effects in early embryogenesis

Maternal and paternal gametophytic effects occur in embryogenesis when the phenotype of the embryo depends on the genotype of the egg or sperm, respectively (reviewed in Armenta-Medina & Gillmor, 2019). We previously used mutations in early-acting *EMBRYO DEFECTIVE (EMB)* genes to test the activity of wt paternal alleles of 49 different *EMB* genes (Del Toro-De León et al., 2014). Because most *emb* mutants are lethal when homozygous, heterozygous (*emb/*+) plants must be used for these assays. For 40 *EMB* genes, we found that wt sperm could not complement embryos derived from mutant *emb* eggs until 3-5 dap, i.e. these 40 *emb* mutants showed a transient maternal effect. For 11 of these 40 *EMB* genes, paternal effects were tested by performing crosses using *emb/*+ plants as fathers. Transient embryo phenotypes were only observed when these *emb/*+ mutants were used as mothers, and not as fathers, indicating that the phenotypes were due to a maternal effect and not to haploinsufficiency (Del Toro-De León et al., 2014).

To extend this analysis, we tested 24 additional *emb* mutants for maternal and paternal effects in early embryogenesis. Phenotypes of embryos derived from hand-selfed Col *emb/*+ plants, from *emb/*+ mothers crossed with wt Col fathers, and from wt Col mothers crossed with *emb/*+ fathers were scored at 2, 3 and 5 dap (Figure 1, Supplemental Table 1, Supplemental Data 1). All 24 *emb* mutants showed maternal effects beginning at 2 or 3 dap, which typically decreased by 5 dap (Figure 1B), and in almost all cases disappeared by 14 dap (Supplemental Table 1). The onset of paternal allele activity (i.e. a decrease in the maternal effect) can be detected as a decrease in the frequency of mutant phenotypes in embryos derived from *emb/*+ x Col crosses (the father contributes two wild type alleles), compared to *emb/*+ x *emb/*+ crosses (the father contributes a single wild type allele). Using this criterion, paternal allele activity could first be detected at 2, 3 or 5 dap, depending on the *EMB* gene (Figure 1D & Supplemental Table 1). To determine if any of the 24 *emb* mutants also had a paternal effect, we used wt Col plants as mothers in crosses with *emb/*+ fathers. Paternal effects were less common than maternal effects (compare Figures 1B&C). *nse1, nse3, miro1, iyo* and *pect1* showed transient paternal effects of the same magnitude as their transient maternal effect (Figure 1D, Supplemental Table 1). *qqt2, gex1, zyg3, gle1* and *fac1* showed transient paternal effects weaker than their transient maternal effects, and the remaining 14 *emb* mutants showed no paternal effect at all (Figure 1C&D & Supplemental Table 1). In summary, maternal and paternal effects were tested for 24 *emb* mutants from this study and for 11 *emb* mutants in Del Toro-De Leon et al. (2014). Maternal effects were seen for all 35 *emb* mutants, a paternal effect equal to the maternal effect was seen for 5 *emb* mutants, and a weak paternal effect was seen for an additional 5 *emb* mutants. These results show that for 30 out of 35 *EMB* genes, the maternal allele plays a more important functional role in early embryogenesis than the paternal allele.

**Figure 1.**
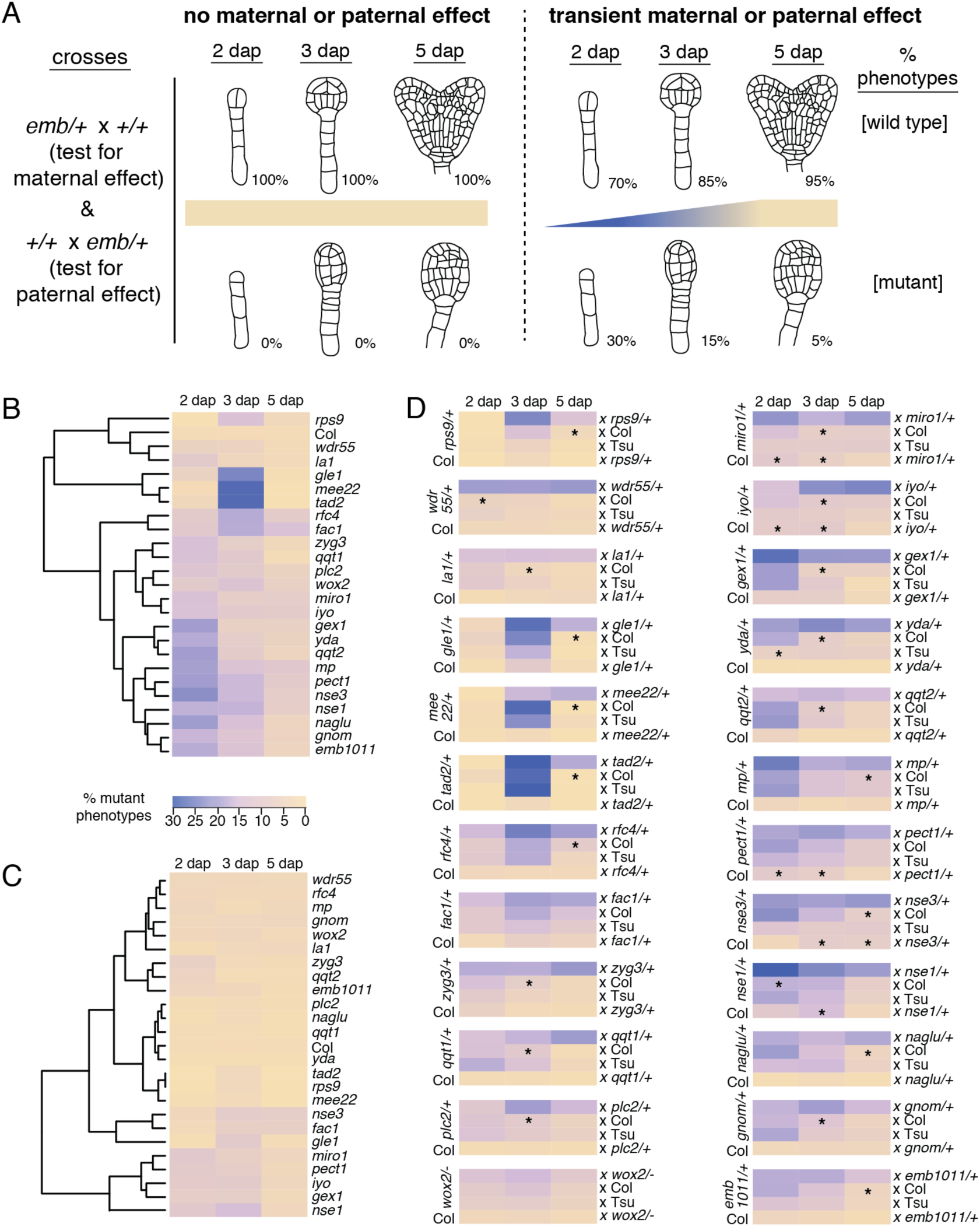
Maternal effects of *emb* mutants are more common than paternal effects, and are similar between isogenic Col and Col x Tsu hybrid embryos. (A) Schematic representations of results for crosses to test maternal and paternal effects of *emb* mutants. Left, percentage of wild type and mutant phenotypes in the absence of maternal and paternal effects. Right, hypothetical example of penetrance of mutant phenotypes when maternal or paternal effects are seen. (B) Maternal effects observed in embryos from *emb/*+ x Col crosses at 2, 3, and 5 dap. (C) Paternal effects observed in embryos from Col x *emb/*+ crosses at 2, 3, and 5 dap. (D) Mutant phenotypes observed in embryos of *emb/*+ x *emb/*+; *emb/*+ x Col; *emb/*+ x Tsu; and Col x *emb/*+ plants at 2, 3, and 5 dap. For *emb/*+ x Col crosses, the asterisk denotes the first stage where a significant decrease in mutant phenotypes could be observed compared with *emb/*+ x *emb/*+ crosses. For *emb/*+ x Tsu crosses, the asterisk marks a significant difference from *emb/*+ x Col. For Col x *emb/*+ crosses, the asterisk denotes stages with both a significant difference with Col x Col control crosses, and no significant difference with *emb/*+ x Col crosses (i.e. equal maternal and paternal effects). Statistical significance p<0.05, determined by Fisher’s two-tailed test. The intensity of purple denotes the % of mutant phenotypes observed in embryos resulting from the indicated crosses. See Supplemental Table 1 and Supplemental Data 1 for full information.

The transient maternal and paternal effects seen for these 35 *emb* mutants could be due to a requirement for gene function in the egg, sperm, zygote, embryo, or a combination of these cell types. To try to distinguish between gametic and zygotic requirements for *EMB* function, we looked for presence of these 35 *EMB* genes in transcriptome datasets for sperm (Borges et al., 2008), eggs, zygotes and embryos (Zhao et al., 2019). 33 *EMB* genes had transcripts in zygotes and embryos, 31 of these 33 genes had egg cell transcripts, and 12 of the 33 *EMB* genes were called as present in the sperm cell transcriptome (Supplemental Table 2). *GEX1* had a massive signal in the sperm cell, strong expression in the egg cell, and progressively decreasing transcripts in the zygote and embryo, strongly suggesting that the paternal and maternal effects observed in early *gex1/*+ embryos are gametophytic (Alandete-Saez et al., 2011). *MONOPTEROS (MP)* and *WUSCHEL HOMEOBOX 2 (WOX2)*, the only genes with zygote and embryo expression but no detected egg cell transcripts, showed maternal effects but no paternal effects, suggesting that the maternal effects for these genes are zygotic maternal effects. The remaining genes had transcripts in the egg, zygote and embryo, making it impossible to distinguish between maternal gametophytic and maternal zygotic effects.

### Preglobular Arabidopsis embryos show a maternal transcript bias

Hybridization of polymorphic ecotypes is necessary to distinguish between RNA sequencing reads originating from the maternal and paternal genomes. However, crossing different ecotypes is known to affect parent-of-origin contributions to early embryos (Del Toro-De León et al., 2014). Notably, transient maternal effects for 11 *emb* mutants were almost identical between isogenic *emb/*+ x Col and hybrid *emb/*+ x Tsu crosses (Del Toro-De León et al., 2014), suggesting that parent-of-origin effects in this hybrid combination are close to the isogenic Col reference strain. To further test this hypothesis, we crossed the 24 additional *emb/*+ mutants from the current study to Tsu, and compared the results with crosses to Col. For 23 of 24 *emb* mutants, transient maternal effects (i.e. *EMB* paternal allele activity) were statistically indistinguishable between *emb/*+ x Col and *emb/*+ x Tsu crosses (Figure 1D & Supplementary Table 1). From the combined results for the 35 *EMB* genes from this study and from Del Toro-De León et al., (2014), we conclude that the functional contributions of paternal alleles are very similar between isogenic Col and the Col x Tsu hybrid.

The similarity of paternal functional contributions between isogenic Col and Col x Tsu, and the large number of genomic DNA polymorphisms between Col and Tsu (Nordborg et al., 2005), make the Col x Tsu hybrid ideal for monitoring parent-of-origin contributions to early embryo transcriptomes. Our analysis of the Col and Tsu ecotypes identified 316,633 variants, of which 161,026 corresponded to exons in 22,741 genes (see Methods). We crossed plants of the Col and Tsu accessions in both directions (Col x Tsu and Tsu x Col), and generated transcriptomes from embryos at the zygote-1 cell stage (zyg-1cell), octant, globular, heart, torpedo, bent and mature stages. These transcriptomes exhibited several different classes of parent-of-origin bias. Genes showing reciprocal maternal bias (RMB) or reciprocal paternal bias (RPB) were biased in both directions of the cross. Genes with reciprocal Col bias (RCB) or reciprocal Tsu bias (RTB) showed higher expression of the Col or Tsu allele, respectively, regardless of the direction of the cross. No reciprocal bias (NRB) genes were those that did not show reciprocal maternal, paternal, Col or Tsu bias. General transcriptome information, complete gene expression data, and complete ASE data are found in Supplemental Data 2. ASE results for the zyg-1cell through heart stages are shown in Figure 2A, and data for all stages are presented in Figure 2B.

**Figure 2.**
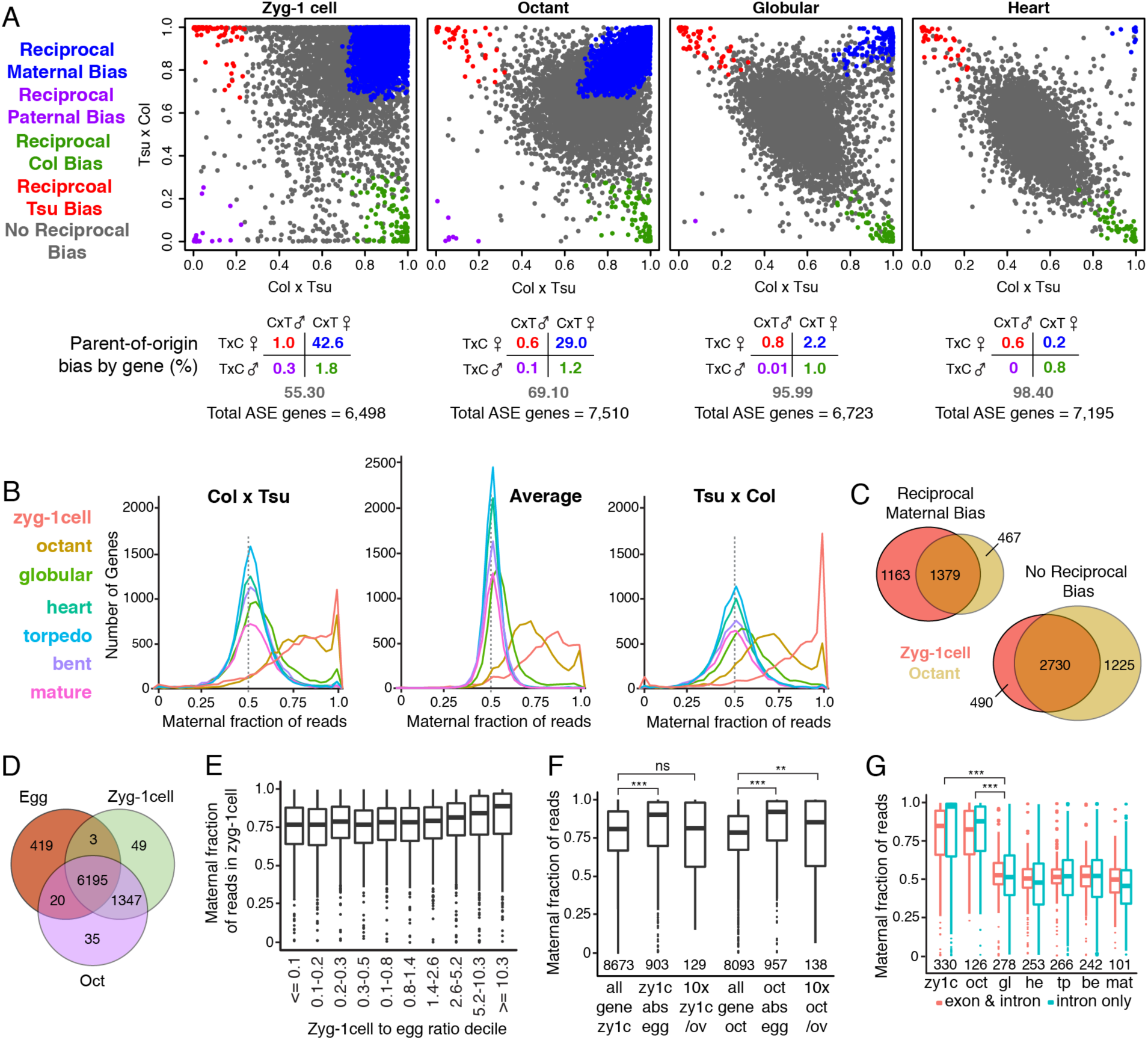
Maternal transcripts predominate in preglobular embryos. (A) Top, maternal fractions of reads for genes in embryos from Col x Tsu and Tsu x Col crosses, at the zyg-1cell, octant, globular and heart stages, where 1.0 is exclusively maternal and 0.0 is exclusively paternal. RMB, reciprocal maternal bias; RPB, reciprocal paternal bias; RCB, reciprocal Col bias; RTB, reciprocal Tsu bias; NRB, no reciprocal bias. Bottom, percentage of genes in each reciprocal bias category, or with no reciprocal bias (grey). (B) Number of genes corresponding to maternal fraction of reads at the zyg-1cell, octant, globular, heart, torpedo, bent and mature stages, in Col x Tsu embryos, Tsu x Col embryos, and the average of the reads for each gene between the two crosses. (C) Venn diagram for the overlap of reciprocal maternal bias and no reciprocal bias genes at the zyg-1cell (orange) and octant (tan) stages. (D) Venn diagram for genes with transcripts present in the egg, zyg-1cell and octant embryo. (E) Maternal fraction of reads for genes at the zyg-1cell stage, based on their zyg-1cell to egg expression ratio, separated into deciles. The Pearson’s value for the correlation of maternal fraction with zyg-1cell to egg ratio is, 0.07 (p=2.7e^-7^) (F) Maternal fraction of reads for all genes expressed in zyg-1cell embryos; genes expressed at the zyg-1cell and absent from egg (*p*=3.03e^-15^); genes expressed ten times higher at the zyg-1cell embryo than in the ovule (*p*=9.31e^-3^); all genes expressed in the octant embryo; genes expressed in the octant embryo and absent from the egg (*p*=2.19e^-22^); and genes expressed ten times higher in the octant embryo compared to the ovule (*p*=1.03e^-02^). (G) Maternal fraction of reads for genes with reads mapping to introns, and reads mapping to exons and introns, at the zyg-1cell through mature stages. Zyg-1cell exon & intron reads vs globular exon & intron reads (*p*=4.9e^-49^). Zyg-1cell intron reads vs globular intron reads (*p*=4.06e^-36^). Octant exon & intron reads vs globular exon & intron reads (*p*=5.63e^-32^). Octant intron reads vs globular intron reads (*p*=1.84e^-24^). *p* values for results of a Wilcox test comparing the maternal fraction of reads between samples are listed in parentheses. ***, p < 0.001; ** p<0.01

Thousands of genes showed reciprocal maternal bias at the zyg-1cell and octant stages (Figure 2A & B; Supplemental Data 2). In zyg-1cell embryos, 42.6% of genes showed RMB and 55.3% showed NRB (n=6,498), while octant stage embryos had 29.0% RMB genes and 69.1% NRB (n=7,510). Many RMB genes were shared between zyg-1cell and octant stage transcriptomes (Figure 2C). At the globular stage, 2.2% of genes were RMB and 95.99% NRB (n=6,723), while heart stage embryos had 0.2% RMB and 98.4% NRB genes (7,195). A small percentage of genes showed paternal bias: 0.3% of genes were RPB at the zyg-1cell stage, 0.1% at the octant stage, and 0.01% at the globular stage. At the zyg-1cell stage, about 3% of genes showed reciprocal Col or Tsu bias, decreasing to less than 2% at later stages (Figure 2A; Supplemental Data 2). From heart to mature stage embryos, ASE results did not significantly vary from one stage to the next (Figure 2B; Supplemental Figure 1).

The maternal bias observed at the zyg-1cell and octant stages could be due to inheritance of transcripts from the egg, preferential transcription of maternal alleles, or both. Comparison of our data with a recent egg cell transcriptome (Zhao et al., 2019) revealed that the majority of gene transcripts in zyg-1cell and octant stage embryos are also found in egg cells: of 7,594 and 7,597 genes represented by zyg-1cell and octant embryo transcripts, 6,198 and 6,215 were present in the egg cell, respectively (Figure 2D; Supplemental Data 3). However, no correlation between the relative level of transcripts in the egg vs zyg-1cell or octant embryo, and maternal transcript bias in zyg-1cell or octant embryos was detected (Figure 2E; Supplemental Figure 2). By contrast, zyg-1cell transcripts absent from the egg (n=903) were 90.2% maternal, versus 80.8% maternal for all zyg-1cell genes; octant stage transcripts absent from the egg (n=957) were 92.0% maternal versus 78.5% for all octant genes (Figure 2F; Supplemental Data 3). The maternal bias of genes with intronic transcripts, indicative of actively transcribed genes (Gaidatzis et al., 2015), was also determined. The median maternal fraction of intronic transcripts was 97.4% in zyg-1cell (n=330 genes) and 87.7% in octant embryos (n=126 genes), while intronic reads at the globular and later stages had no maternal bias (Figure 2G; Supplemental Data 3). These results strongly suggest that the maternal transcript bias seen in zygote and octant stage embryos is at least partially due to preferential transcription of maternal alleles, and are entirely consistent with functional genetic experiments demonstrating predominantly maternal effects for *emb* mutants (Figure 1).

Embryos are surrounded by the maternally-derived seed coat, and thus contamination by maternal tissue needs to be considered when determining parent-of-origin bias in early embryo transcriptomes. An algorithm designed to identify genes enriched in embryo, endosperm and seed coat tissues detected the presence of both embryo and seed coat enriched transcripts in our zyg-1cell and octant embryo datasets (Supplemental Figure 3) (Schon & Nodine, 2017). To test if the maternal bias we observed in our zyg-1cell and octant transcriptomes was influenced by maternal sporophytic transcripts, we used ovule transcriptomes to calculate zyg-1cell/ovule and octant/ovule expression ratios on a gene-by-gene basis. Transcripts for genes enriched ten times in zyg-1cell embryos compared to the ovule (n=129) were 81.3% maternal, compared to a median maternal ratio of 80.8% for all zygotic genes (n=8673). Transcripts for genes enriched ten times in octant embryos compared to the ovule (n=138) were 85.2% maternal, compared to a median 78.5% maternal bias for all octant embryo transcripts (n=8093) (Figure 2F & Supplemental Data 3). We also compared the maternal bias for all genes at the zyg-1cell and octant stages with their embryo/ovule expression ratio. No correlation was found between maternal bias and zyg-1cell/ovule or octant/ovule expression ratio (Supplemental Figure 4 & Supplemental Data 3). These results demonstrate that genes with higher expression in the ovule than the embryo show no more maternal bias than genes which are higher in the embryo than the ovule. We conclude that contamination by seed coat transcripts does not have a significant effect on the overall maternal transcript bias observed in zyg-1cell and octant embryos.

To further validate the transcriptome results above, we checked for correlations between transcriptome data and functional or reporter gene data. *emb* mutants with both maternal and paternal effects are likely to be haploinsuficient in early embryogenesis, and would be expected to have similar levels of maternal and paternal transcripts, while *emb* mutants with only maternal effects must have maternal expression and may or may not have similar levels of paternal expression. Of the 35 *EMB* genes with functional data from this study or Del Toro-De León et al. (2014), 19 are represented by parent-of-origin data in the Col x Tsu transcriptome (Supplemental Table 2). Of these 19 *EMB* genes, the five *emb* mutants with both maternal and paternal effects (*iyo, nse3, qqt2, fac1* and *gle1*) have similar amounts of maternal and paternal reads, while the 14 *emb* mutants with only maternal effects have maternal reads with varying ratios of paternal reads (Supplemental Table 2). Parent-of-origin expression for three reciprocally maternally biased genes is also consistent with reporter gene data. *NUCLEAR FACTOR Y SUBUNIT B2 (NF-YB2)* showed almost exclusively maternal transcripts at the zyg-1cell and octant stages, and strong maternal transcript bias at the globular stage, in agreement with strictly maternal expression of an *pNF-YB2∷GUS* marker (Siefers et al., 2009) at 2 and 3 dap (Figure 3 & Supplemental Table 3). *GRAND CENTRAL* (*GCT)* and *CENTER CITY (CCT)* also showed reciprocal maternal transcript bias at the zyg-1cell and octant stages, in agreement with *gGCT-GUS* and *pCCT∷GUS* reporters which showed maternally biased GUS marker expression at 2 and 3 dap (Del Toro-De León et al., 2014; Supplemental Table 3). The correlation between transcriptome, functional and marker data is consistent with faithful representation of parent-of-origin biases in the transcriptome reported here.

**Figure 3.**
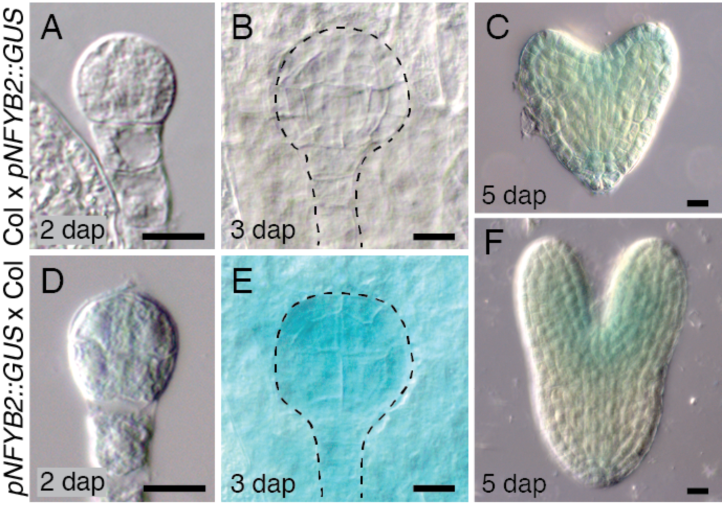
Maternally biased expression of *pNF-YB2∷GUS* marker. Expression of *pNF-YB2∷GUS* from paternal (A-C) and maternal (D-F) alleles is shown at 2, 3 and 5 dap. See Supplemental Table 3 for quantification of GUS expression. Scale bar = 10µm.

In contrast to the maternal transcript bias in Col x Tsu embryos at the zyg-1cell and octant stage, equal maternal and paternal transcripts were found in 1-2 cell and octant Col x Cvi embryos (Nodine & Bartel, 2012), and Col x L*er* zygotes (Zhao et al., 2019). The Col x Tsu, Col x Cvi, and Col x L*er* hybrid combinations were previously found to be divergent for paternal allele activation (Del Toro-De León et al., 2014), suggesting that the different parent-of-origin transcriptome contributions found in these three hybrids reflect biological differences between the ecotypes used. For example, the equal maternal and paternal transcripts in Col x Cvi may be due to the unique chromatin status of the Cvi ecotype, which has genome-wide hypomethylation at CG sites (Schmitz et al., 2013; Pignatta et al., 2014), and more open chromatin (Tessadori et al., 2009), leading to an increase in transcription in Col x Cvi hybrids (Baroux et al., 2013). The predominance of maternal effects observed in analysis of *emb* mutants in Col and Col x Tsu embryos, and the maternal transcript bias observed in Col x Tsu embryos, may also have an epigenetic basis. The Arabidopsis egg cell lacks expression of the CG maintenance methyltransferase *MET1* (Jullien et al., 2012), and has low levels of CG methylation, likely due to hemi-methylation of CG sites (Ingouff et al., 2017). By contrast, the Arabidopsis sperm cell has steady levels of CG methylation (Ingouff et al., 2017). When these parental genomes are united in the zygote, the resulting asymmetry in CG methylation could result in greater transcriptional silencing of the paternal genome than the maternal genome, due to silencing proteins like SU(VAR)3-9 HOMOLOG2 (SUVH2) and METHYL BINDING DOMAIN 5 (MBD5), MBD6 & MBD7 that specifically bind to symmetrically methylated CpG sites to (Johnson et al., 2008; Zemach & Grafi, 2003). The variation in CG methylation that exists throughout the genome could explain why parent-of-origin bias varies between genes, and might underlie some of the differences observed between ecotypes, given the variation in DNA methylation that exists between Cvi and other accessions (Schmitz et al., 2013; Pignatta et al., 2014). A recent study in rice found large differences in siRNA populations of eggs and sperm (Li et al., 2020), which could influence transcription of maternal and paternal genomes in rice zygotes (Anderson et al., 2017).

Changes in mRNA metabolism between ecotypes or hybrids could also explain differences in transcript populations in early embryos. A transcriptome of L*er* x Col embryos that used random primers to generate libraries for sequencing found a maternal bias in early embryos (Autran et al., 2011). Significant differences between total mRNA and polyA mRNA populations are known in animal zygotes (Aanes et al., 2011), have previously been found in maize (Grimanelli et al., 2005), and are very likely to exist in Arabidopsis as well. For example, mRNAs for *WOX2*, one of the most important transcription factors in early embryogenesis in Arabidopsis, have been detected in egg cells by *in situ* hybridization (Haecker et al., 2004), but are absent from an egg cell transcriptome produced using oligodT primers (Supplemental Table 2; Zhao et al., 2019), suggesting that the increase in *WOX2* transcripts detected by RNAseq after fertilization is at least partly due to polyadenylation of existing *WOX2* mRNAs.

## Conclusion

The genetic and transcriptome data reported here show a predominant role for the maternal genome during early Arabidopsis embryogenesis, and point to Col x Tsu as an opportune hybrid for studies of parent-of-origin gene regulation in Arabidopsis. Why might parent-of-origin transcript ratios vary between different hybrids? The parental conflict theory would predict maternal silencing of paternal alleles to be adaptive (Baroux & Grossniklaus, 2015). However, in an inbred species like *Arabidopsis thaliana*, relative changes in early maternal and paternal transcription may have less functional relevance than they would in an outcrossing species, allowing for rapid evolution in parent-of-origin gene expression in seeds of different ecotypes (Geist et al., 2019). For example, the RNA-dependent DNA methylation pathway, as well as histone methylation and histone replacement, have been demonstrated to regulate expression of paternal alleles in early Arabidopsis embryos (Autran et al., 2011). Yet loss of function phenotypes of these pathways have only subtle phenotypes during embryogenesis (Pillot et al., 2010; Autran et al., 2011). By contrast, loss of the maternal RdDM pathway in the outcrossing species *Brassica rapa* causes severe phenotypes during reproduction (Grover et al., 2018), demonstrating the importance of maternal RdDM regulation of seed development in a close relative of Arabidopsis. A better understanding of the mechanisms that regulate maternal and paternal transcript ratios in the zygote and early embryo will reveal more about the biological role that maternal dominance plays in early embryogenesis in *Arabidopsis thaliana*, and will facilitate comparative studies of maternal and paternal regulation of embryogenesis in species of agronomic importance like rice and maize.

## Methods

In notation of crosses, the mother is always listed first. *emb/*+ crosses and reporter gene studies (NF-YB2 reporter ABRC stock CS67020) were conducted as in Del Toro-De León et al. (2014). To produce hybrid embryos for ASE analysis, the inflorescence of the female parent was pruned to leave only the two oldest flower buds, which were pollinated 4 hours after emasculation. Embryos from reciprocal crosses of Col-0 (ABRC stock CS60000) and Tsu-1 (ABRC stock CS6926) were collected and processed as in Xiang et al. (2011). Zygote and octant stage embryos were washed in the dissection solution by moving away from ovule and endosperm tissue and then pipetting into eppendorf tubes on dry ice. Globular and later stage embryos were washed using several changes of dissection solution. ∼300 zygote and 1-cell stage embryos (referred to as zyg-1cell stage) were collected at 32 hours after pollination (32 hap), ∼200 octant embryos at 60 hap, ∼100 globular embryos at 72 hap, and ∼100 heart, ∼60 torpedo, ∼30 bent and ∼30 mature embryos, based on morphological stage, as well as ∼10 unfertilized Col ovules. Two biological replicates were collected for all stages in both directions, except for Col x Tsu zyg-1cell stage (one biological replicate). Raw data from Tsu-1 genome re-sequencing (Ossowski et al., 2008) was used to call variants regarding Col-0 (TAIR 10). Reads were trimmed from both ends to have a mean phred quality of at least 30 in 4bp windows using fastp V 0.19.6 (Chen et al., 2018); they were then aligned against Col-0 genome (TAIR 10) using the Burrows-Wheeler aligner V 0.7.17-r1188 (Li & Durbin, 2009) with the aln algorithm, allowing up to 10% of read length as edit distance (-n 0.1); after sorting and marking duplicates with sambamba V 0.6.8 (Tarasov et al., 2015), potential variants were detected using freebayes V 1.0.0 (Garrison and Marth, 2012). Variants were called if they had a minimum phred quality of 20 and a maximum of 0.05 normalized likelihood of being heterozygous. Diploid genome and annotation for the hybrid were generated by substituting these variants into the TAIR10 genome assembly and Araport 11 annotation (Cheng et al., 2017).

For quantification of transcript abundance, reads were aligned against the diploid genome using STAR V 2.7.0d (Dobin et al., 2013), with the following flags: --alignIntronMax 900 -- alignMatesGapMax 900. Reads aligned to genes were then counted using featureCounts V 1.6.0 (Liao et al., 2014) with the following flags: -M -p -O –fraction. Counts for both alleles were summed for each gene, and converted to reads per million (RPM). Abundance ratios between samples were calculated only for genes with at least 20 RPM in at least one of the samples and after adding 5 RPM to the abundance in each tissue, to avoid infinite ratios, and to stabilize the ratios in low-abundance transcripts. To analyze the overlap of present transcripts between stages, genes with more than 10 RPM were considered present and genes with less than 1 RPM were considered absent; genes with transcript levels between 1 and 10 RPM at any stage were ignored, to minimize false call rate. For all comparisons between Col x Tsu embryos and the isogenic Col-0 egg or ovule, only ASE data for the Col x Tsu cross was used, to avoid confounding effects from Tsu maternal transcripts of dissimilar abundance between ecotypes.

For quantification of allele-specific transcript abundance, STAR was run again as previously described, plus the following flags: --outFilterMultimapNmax 1 --outFilterMultimapScoreRange 0. Then, only reads aligned to called variants in genes were counted; a reduced annotation which included only the variable regions of each gene was used, and featureCounts was run as above but without the -M flag; this produced a table of counts for each allele for each gene. For the statistical analysis of allele-specific differential abundance we used the edgeR package (Robinson et al., 2010). Only genes with at least 3 RPM aligned to variants in at least 2 samples were considered for the differential expression analysis. The sum of counts for both alleles was used to calculate library sizes and normalization factors. Tagwise dispersion and the likelihood ratio test were used to obtain FDR-adjusted P values and fold change estimates, which were converted to allelic ratio estimates. For all representations of allelic ratios, all genes with at least 20 reads aligned to variants, considering the sum across replicates, were used. A cutoff of FDR<= 0.05 was used to call significant allelic biases. For analysis of intronic reads, reads were counted only if they aligned to a variant in an intron that did not overlap with any exon of any isoform in the Araport 11 annotation. To diminish the impact of unknown isoforms from the Tsu accession, only reads for the Col x Tsu cross were used. Technical replicates were summed at the count level and treated as a single sample. Raw sequence data are archived in the GEO database (GSE132449).

## Acknowledgements

Seed of *emb* lines and the *pNF-YB2∷eGFP-GUS* reporter were obtained from the Arabidopsis Biological Resource Center. *wox2-1* seed were obtained from Thomas Laux. We are grateful to David Meinke and colleagues for their years of effort assembling and maintaining the *emb* mutant collection, which served as a foundation of this work. Thanks to Wolfgang Lukowitz, Luis Herrera-Estrella, Karina Orozco-Natividad, Daphné Autran and Daniel Grimanelli for comments and suggestions. This research was supported by NRC-GHI and ACRD grants (RD), and by CONACyT Ciencia Básica grant 237480, SEP-CINVESTAV grant 173, and CINVESTAV institutional funds (CSG).

---

**Supplemental Figure 1.**
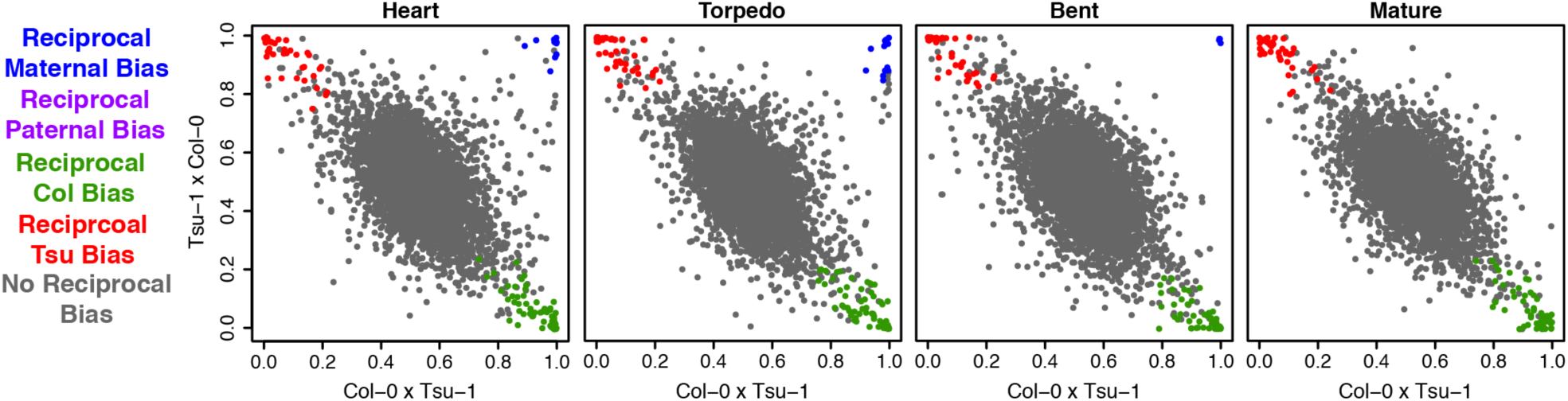
Maternal fraction of reads for genes at the heart, torpedo, bent and mature stages. Maternal fraction of reads for genes in embryos from Col x Tsu and Tsu x Col crosses, where 1.0 is exclusively maternal and 0.0 is exclusively paternal. RMB, reciprocal maternal bias; RPB, reciprocal paternal bias; RCB, reciprocal Col bias; RTB, reciprocal Tsu bias; NRB, no reciprocal bias.

**Supplemental Figure 2.**
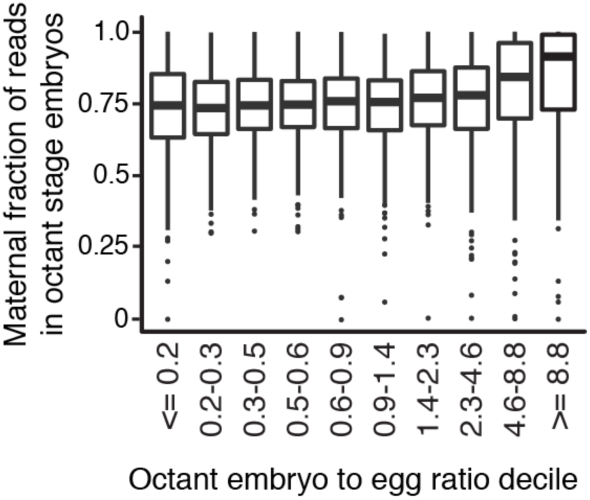
Maternal fraction of reads for genes compared to their octant embryo to egg expression ratios. Maternal fraction of reads in the octant embryo for genes based on their octant embryo to egg expression ratio, separated into deciles. The Pearson’s value for the correlation of maternal fraction with zygote to egg ratio is 0.08 (p=6.7e^-10^).

**Supplemental Figure 3.**
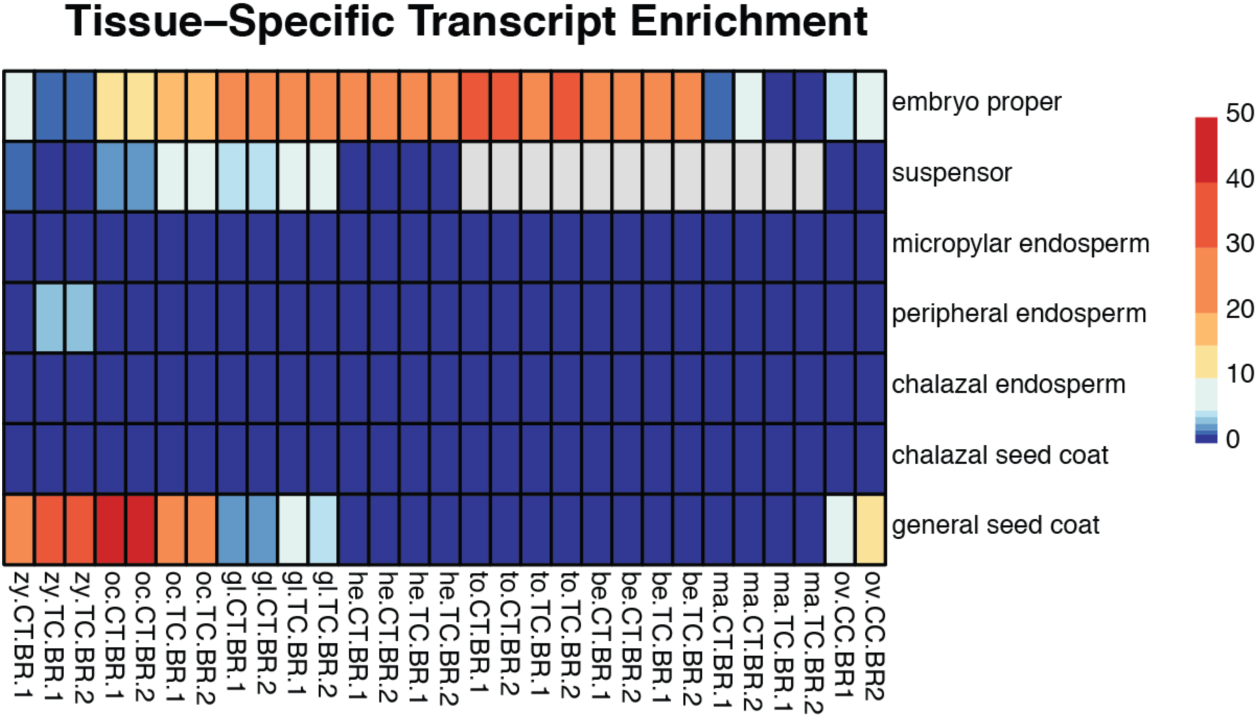
Results of Nodine and Schon tissue enrichment test. Total mapped reads (without regard to parent-or-origin) were run on https://github.com/Gregor-Mendel-Institute/tissue-enrichment-test.

**Supplemental Figure 4.**
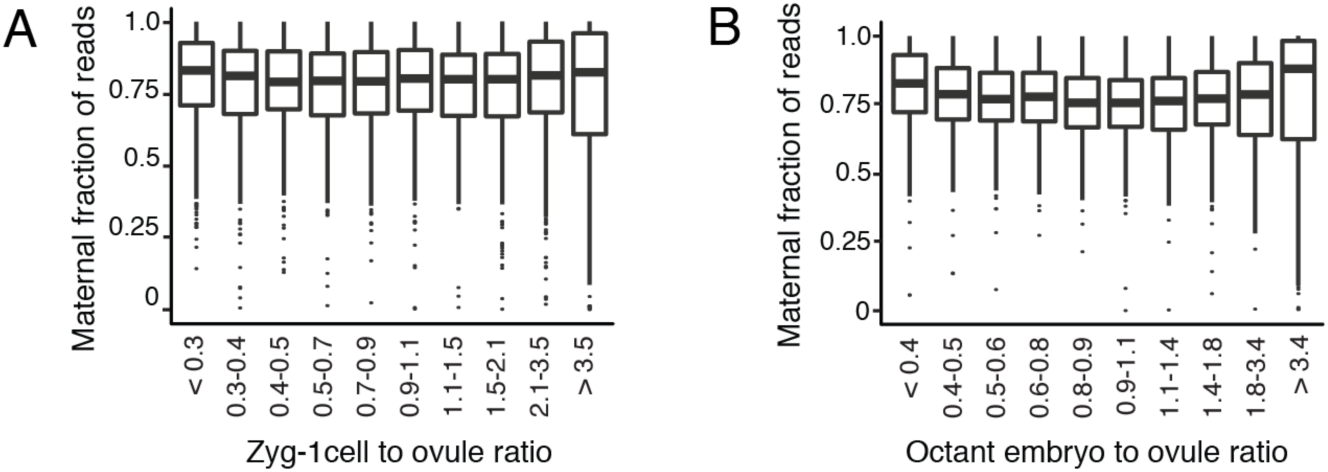
Maternal fraction of reads for genes based on their zyg-1cell and octant stage embryo to ovule expression ratio. (A) Maternal fraction of reads for genes classified into deciles by the ratio of their zyg-1cell to ovule expression (Pearson’s correlation = 0.01, *p*=0.48) (B) Maternal fraction of reads for genes classified into deciles by the ratio of their octant stage embryo to ovule expression (Pearson’s correlation = 0.03, *p*=0.02).

**Supplemental Table 1.**
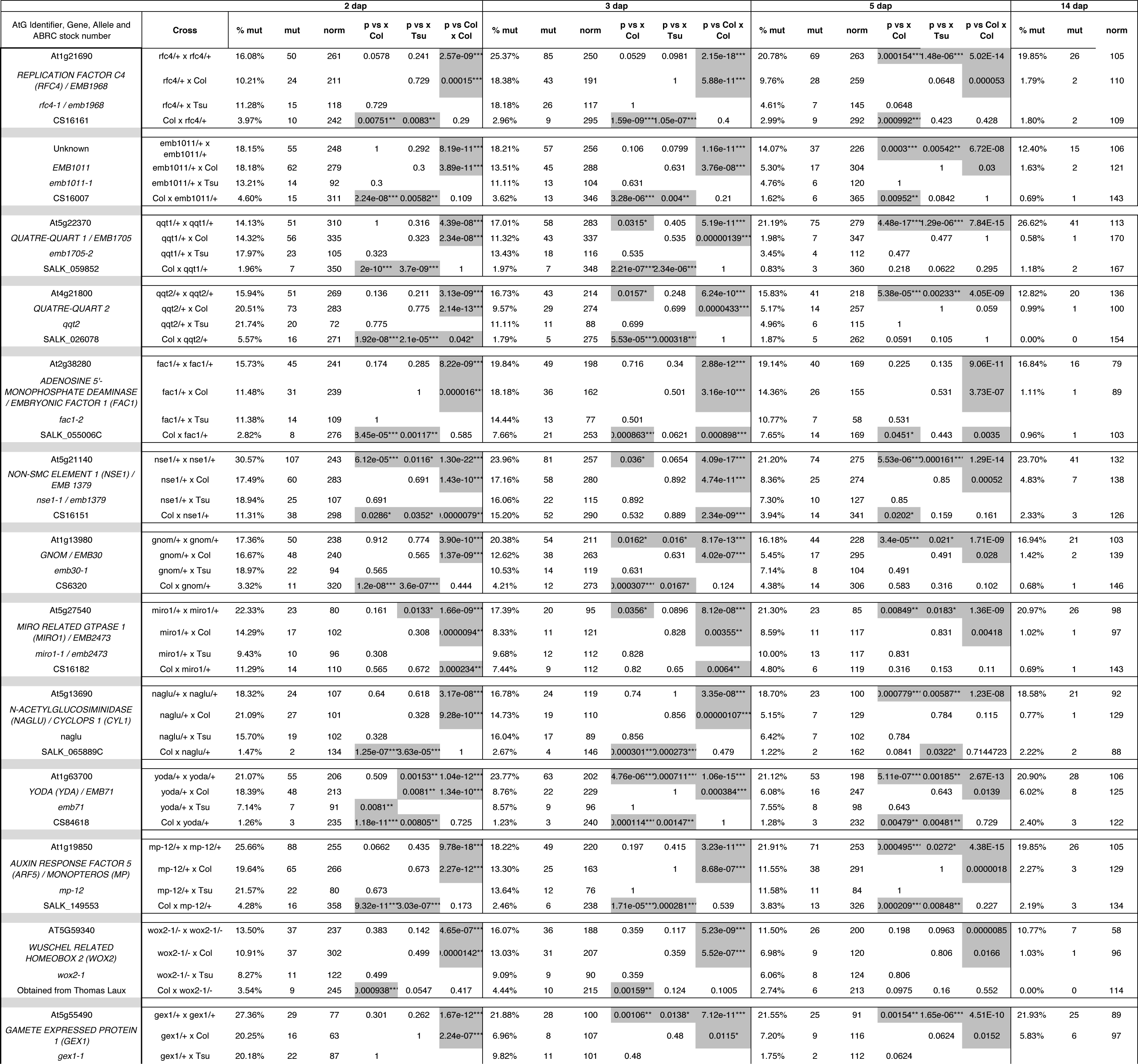
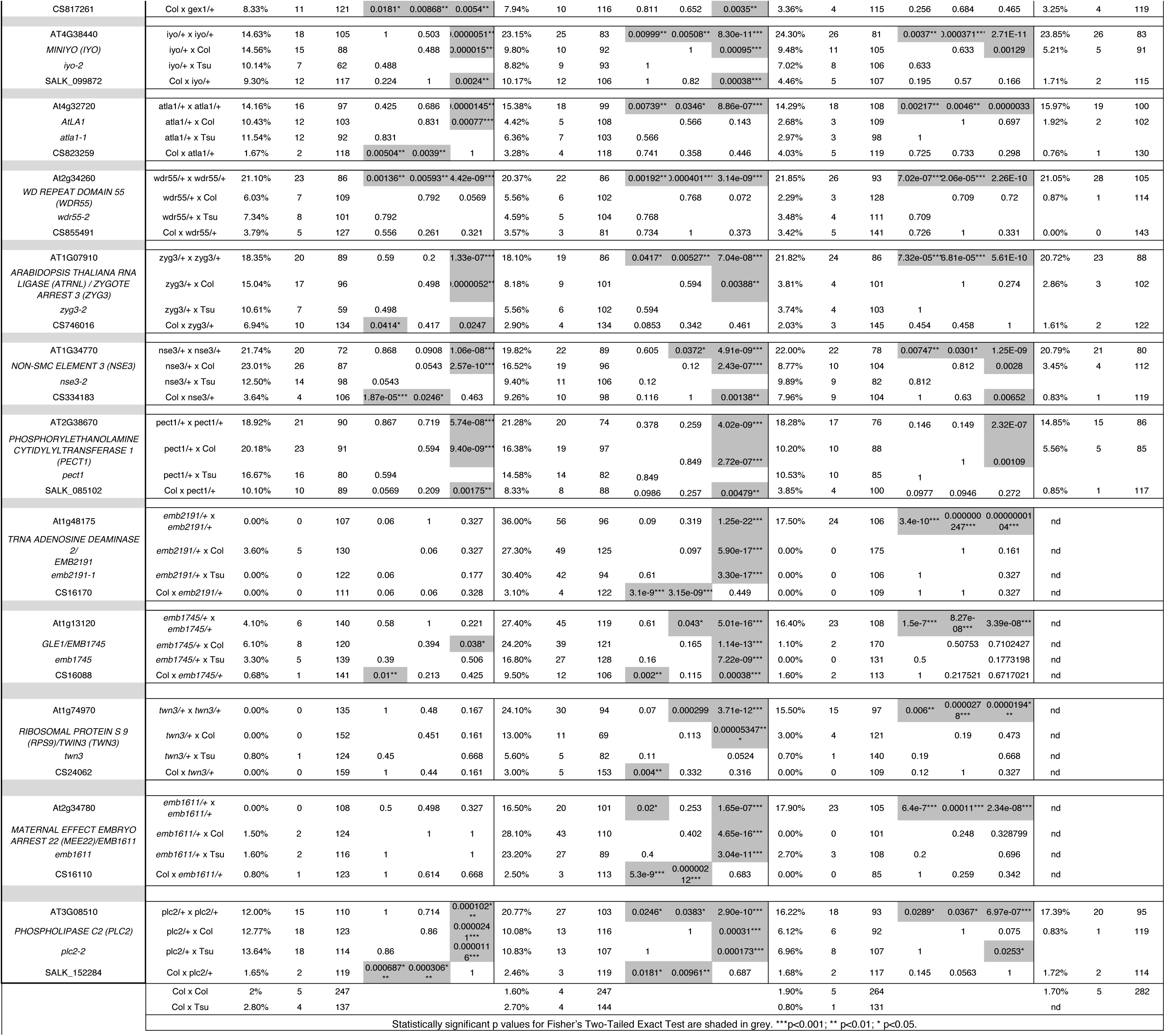
Functional genetic analysis of transient maternal and paternal effects in early Arabidopsis embryogenesis. For each *embryo defective (emb)* mutant, the results of hand self pollinations, crosses to wild type Columbia (Col-0) (CS1092) and Tsu (Tsu-1) (CS28782), are shown. Maternal genotype in crosses is listed first. *emb/*+ (heterozygous) plants were used in all crosses, except for *wox2*, which is homozygous viable and was used as a homozygote. Results are the sum of at least three different siliques from each of two plants. Statistically significant p values for Fisher’s Two-Tailed Exact Test are shaded in grey. ***p<0.001; ** p<0.01; * p<0.05.

**Supplemental Table 2.**
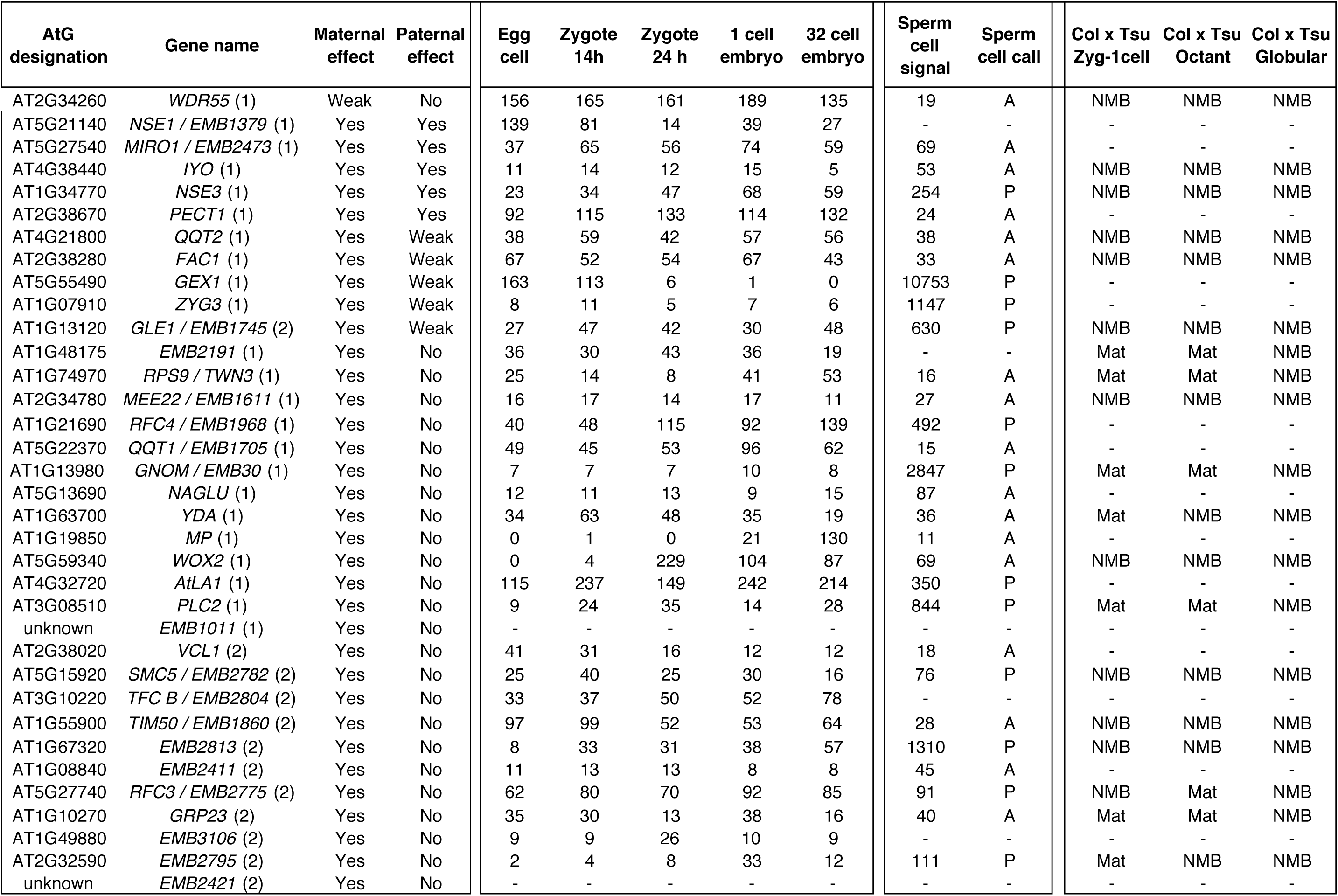
Comparison of transient maternal and paternal effects with egg, zygote and embryo transcriptome data. Maternal and paternal effect data are from this study (1) and from Del Toro-De León et al., 2014 (2). Egg, zygote and embryo expression data (FPKM) from isogenic Col are from Zhao et al., 2019. Sperm cell expression data (ATH1 microarray) and presence (P) absence (A) calls from isogenic Col are from Borges et al., 2008. Col x Tsu hybrid transcriptome data are from this study, and represent data from the Col x Tsu cross only, to make the data most directly comparable with egg and zygote transcriptome data in the Col ecotype, and functional analysis in the Col ecotype. NMB = no maternal bias (maternal fraction adjusted p value > 0.05); Mat = maternal bias (maternal fraction adjusted p value < 0.05).

**Supplemental Table 3.** Comparison of maternal transcript ratios and reporter gene expression for the reciprocal maternal bias genes *NUCLEAR FACTOR Y SUBUNIT B2 (NF-YB2), GRAND CENTRAL (GCT)*, and *CENTER CITY (CCT)*. Data for Col x Tsu (CxT) and Tsu x Col (TxC) transcriptomes at the zyg-1cell, octant globular and heart stages are from Supplemental Dataset 1; p value indicates probability of maternal bias. GUS reporter data shown as fraction of embryos with GUS expression for pNF-YB2∷eGFP/GUS line (ABRC germplasm CS67020) (this study), and pCCT∷GUS and gGCT-GUS (Del Toro-De León et al., 2014) at 1, 2, 3, and 5 dap; p value for Fisher’s Two Tailed Test indicates whether maternal and paternal GUS expression ratios are different from each other. Statistically significant maternal ratios are highlighted in grey.

| Gene ID | Gene name | Zyg-1cell |  |  |  | Octant |  |  |  | Globular |  |  |  | Heart |  |  |  | 2 dap |  |  |  |  | 3 dap |  |  |  |  | 5 dap |  |  |  |  |
| --- | --- | --- | --- | --- | --- | --- | --- | --- | --- | --- | --- | --- | --- | --- | --- | --- | --- | --- | --- | --- | --- | --- | --- | --- | --- | --- | --- | --- | --- | --- | --- | --- |
|  |  | CxT | p | TxC | p | CxT | p | TxC | p | CxT | p | TxC | p | CxT | p | TxC | p | GUS x Col | n | Col x GUS | n | p | GUS x Col | n | Col x GUS | n | p | GUS x Col | n | Col x GUS | n | p |
| At5g47640 | NF-YB2 | 0.974 | 1E-09 | 0.991 | 3E-24 | 0.971 | 5E-16 | 0.92 | 2E-09 | 0.782 | 0.004 | 0.828 | 0.004 | 0.672 | 0.467 | 0.425 | 0.964 | 0.77 | 53 | 0 | 129 | 2E-30 | 0.89 | 91 | 0 | 137 | 4E-51 | 0.76 | 140 | 0.71 | 137 | 0.41 |
| At1g55325 | GCT | 0.897 | 6E-05 | 0.904 | 2E-08 | 0.905 | 4E-07 | 0.826 | 6E-04 | 0.577 | 0.769 | 0.712 | 0.322 | 0.543 | 1 | 0.522 | 1 | 0.84 | 237 | 0 | 118 | 6E-61 | 0.77 | 121 | 0.61 | 271 | 0.002 | 1 | 99 | 1 | 72 | 1 |
| At4g00450 | CCT | 0.778 | 0.017 | 0.716 | 0.013 | 0.832 | 9E-05 | 0.713 | 0.036 | 0.622 | 0.511 | 0.451 | 0.948 | 0.51 | 1 | 0.581 | 0.93 | 0.92 | 166 | 0 | 133 | 1E-70 | 0.82 | 147 | 0.56 | 293 | 2E-08 | 0.99 | 106 | 1 | 99 | 0.38 |

