## Supplemental Data 1 for "Maternal genome dominance in early plant embryogenesis"

**Supplemental Data 1** Phenotypes scored for *embryo defective (emb)* mutants used to generate the data in Figure 1 and Supplementary Table 1. Mutant phenotypes scored are shown for hand self-crosses (*emb/+* x *emb/+*), as well as for crosses to determine maternal effects (*emb/+* x Col) and paternal effects (Col x *emb/+*). Mutant phenotypes scored in maternal and paternal effect crosses to Tsu were the same as those observed in crosses to Col.

### *AtLA1*

## At4g32720

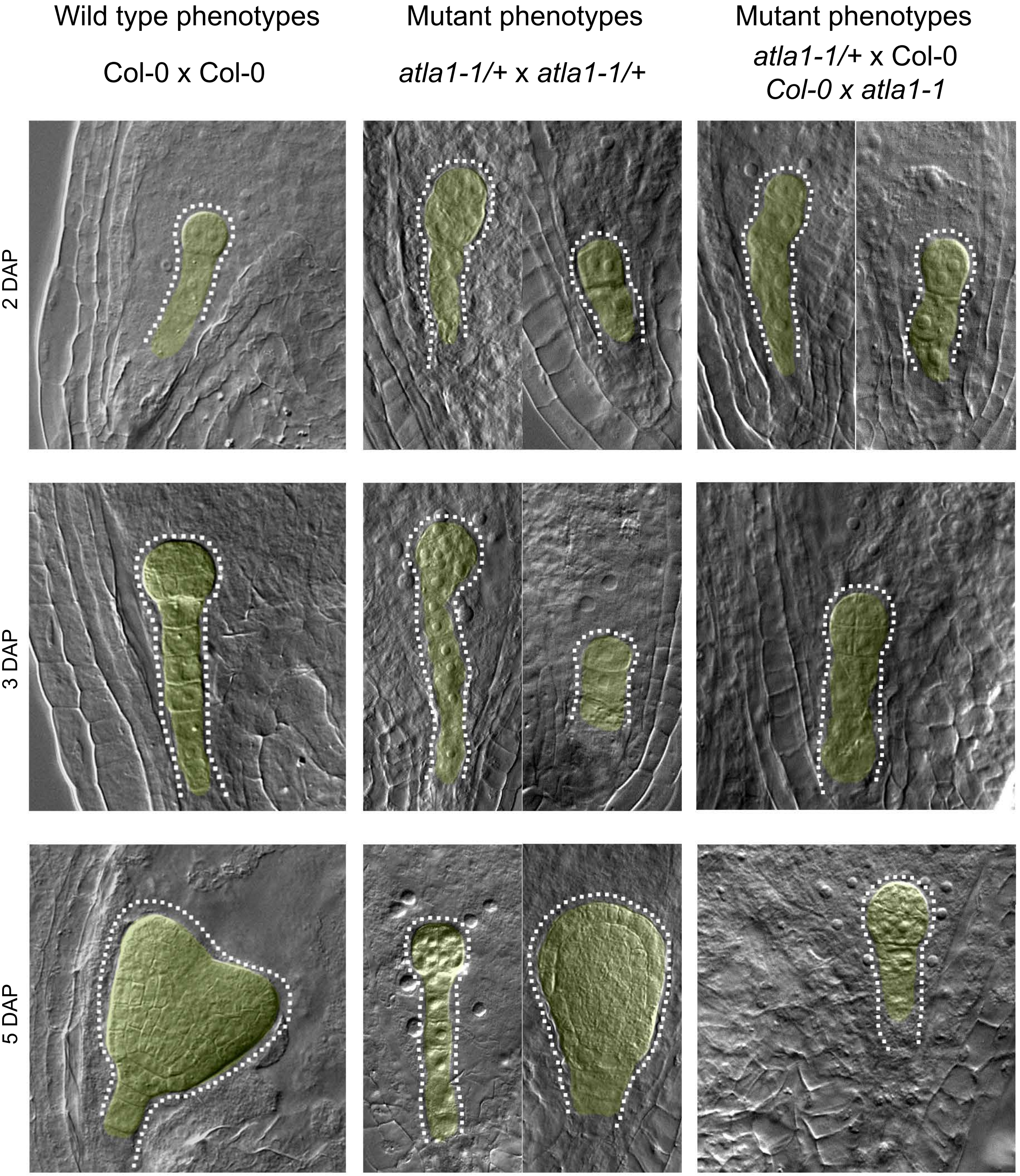


**Protein predicted function:** Encodes AtLa1, a member of the highly abundant phosphoprotein La proteins.

**Seed stock number:** CS823259

### *EMB1011*

## At1g55540

Wilt type phenotypes

Col-0 x Col-0

Mutant phenotypes

*emb1011-1/+* x *emb1011-1/+*

Mutant phenotypes

*emb1011-1/+* x Col-0

Col-0 x *emb1011-1*

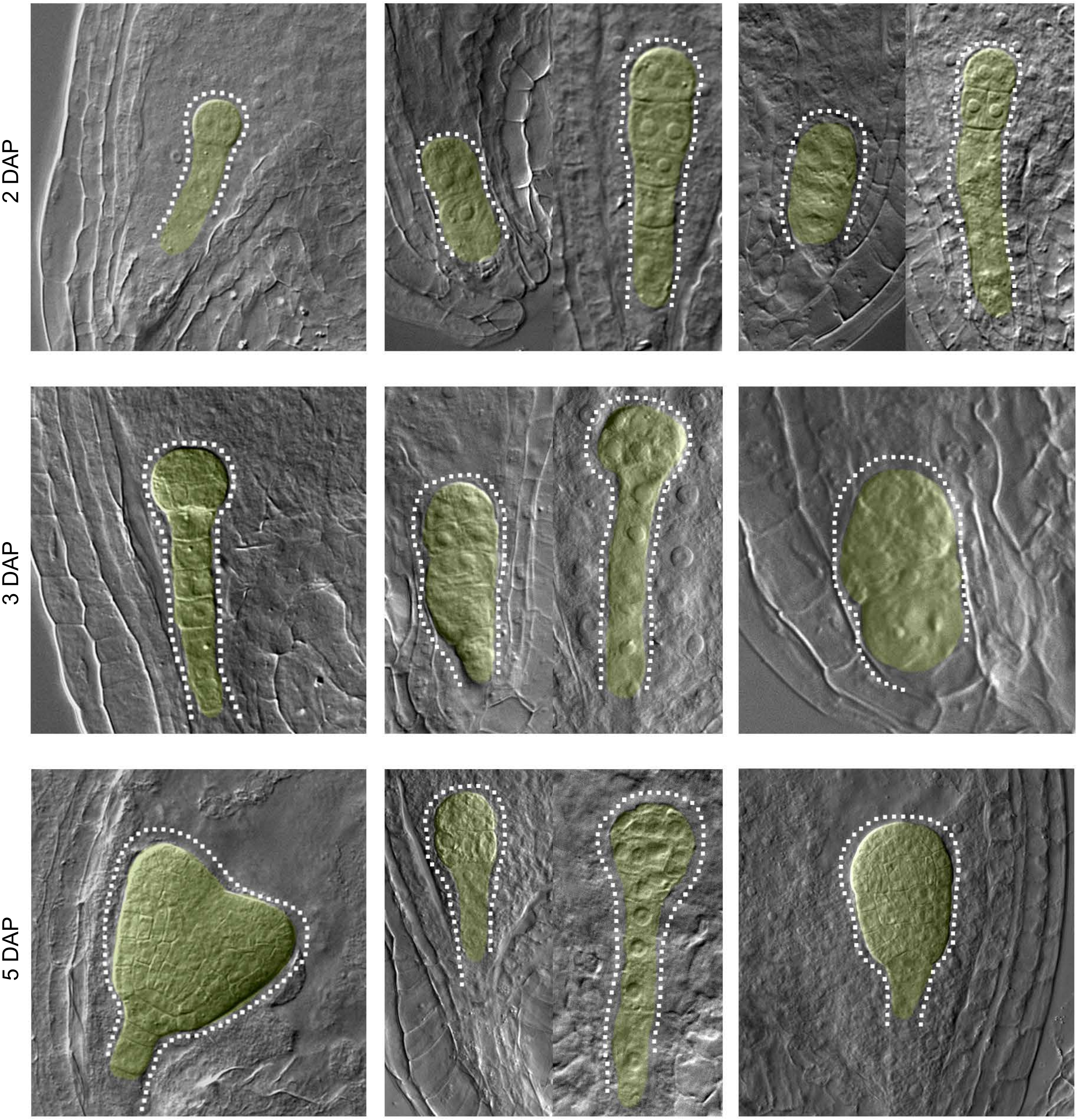


**Protein predicted function:** Nucleoporin protein containing phenylalanine- glycine repeat

**Seed stock number:** CS16007

### *FAC1*

## At2g38280

| Wild type phenotypes | Mutant phenotypes | Mutant phenotypes |
|---|---|---|
| Col-0 x Col-0 | *fac1-2/+* x *fac1-2/+* | *fac1-2/+* x Col-0<br>Col-0 x *fac1-2* |

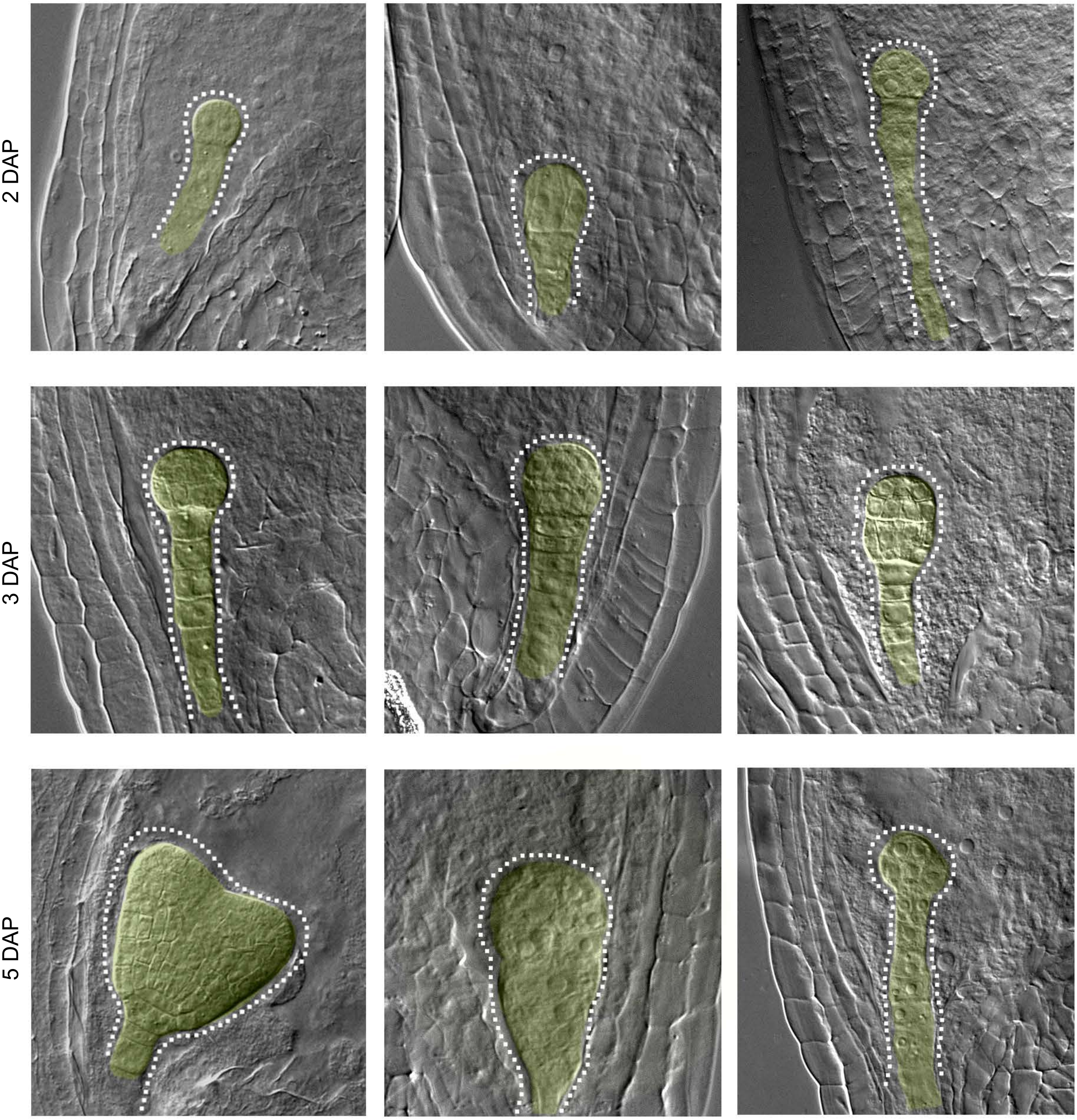


**Protein predicted function:** Encodes a AMP deaminase (AMPD). crucial role for FAC1 in early embryo development.

**Seed stock number:** SALK_055006C

### *GEX1*

## At5g55490

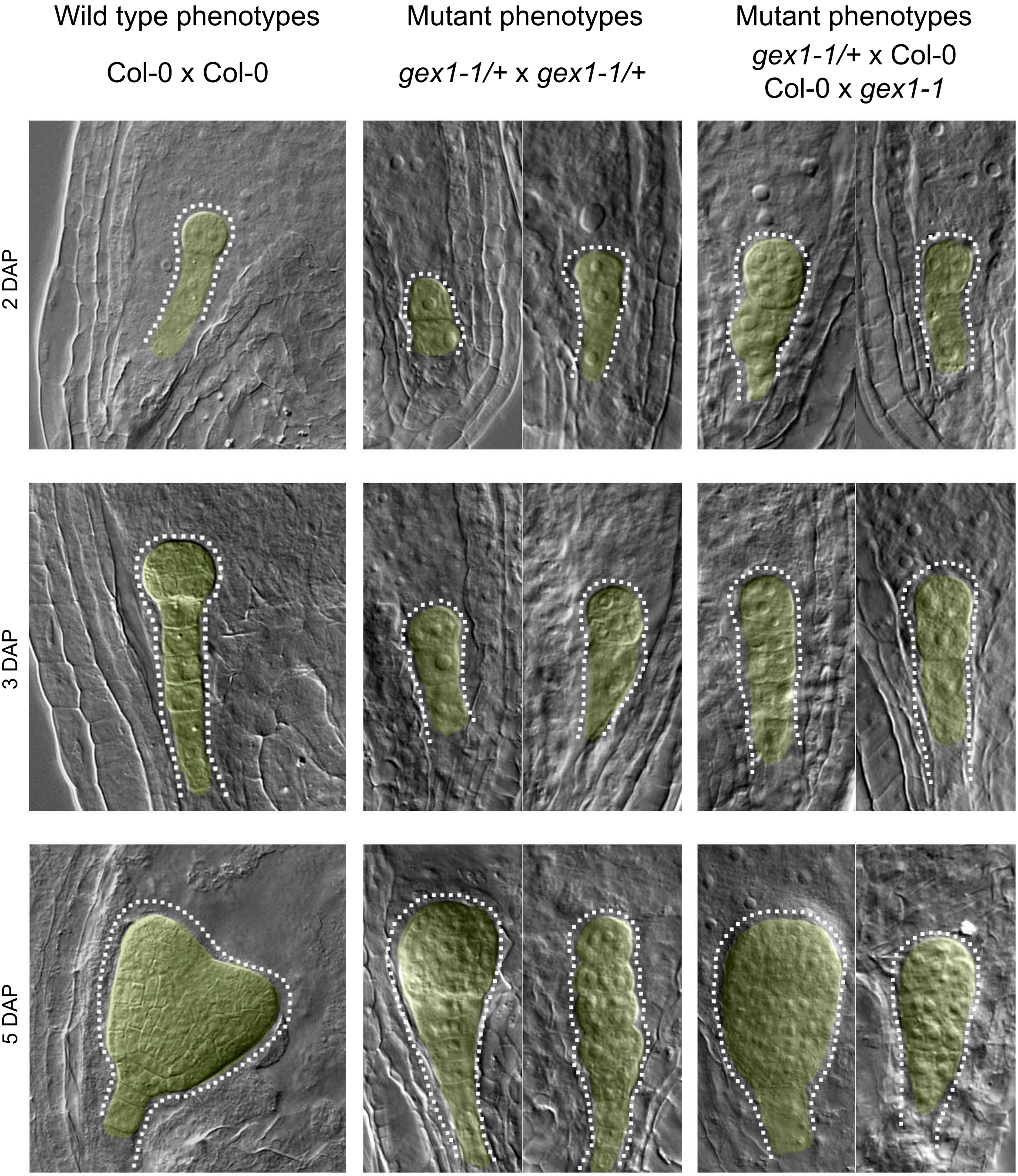


**Protein predicted function:** Encodes a transmembrane domain containing protein that is expressed in pollen germ cells.

**Seed stock number:** CS817261

### *EMB30 / GNOM*

## At1g13980

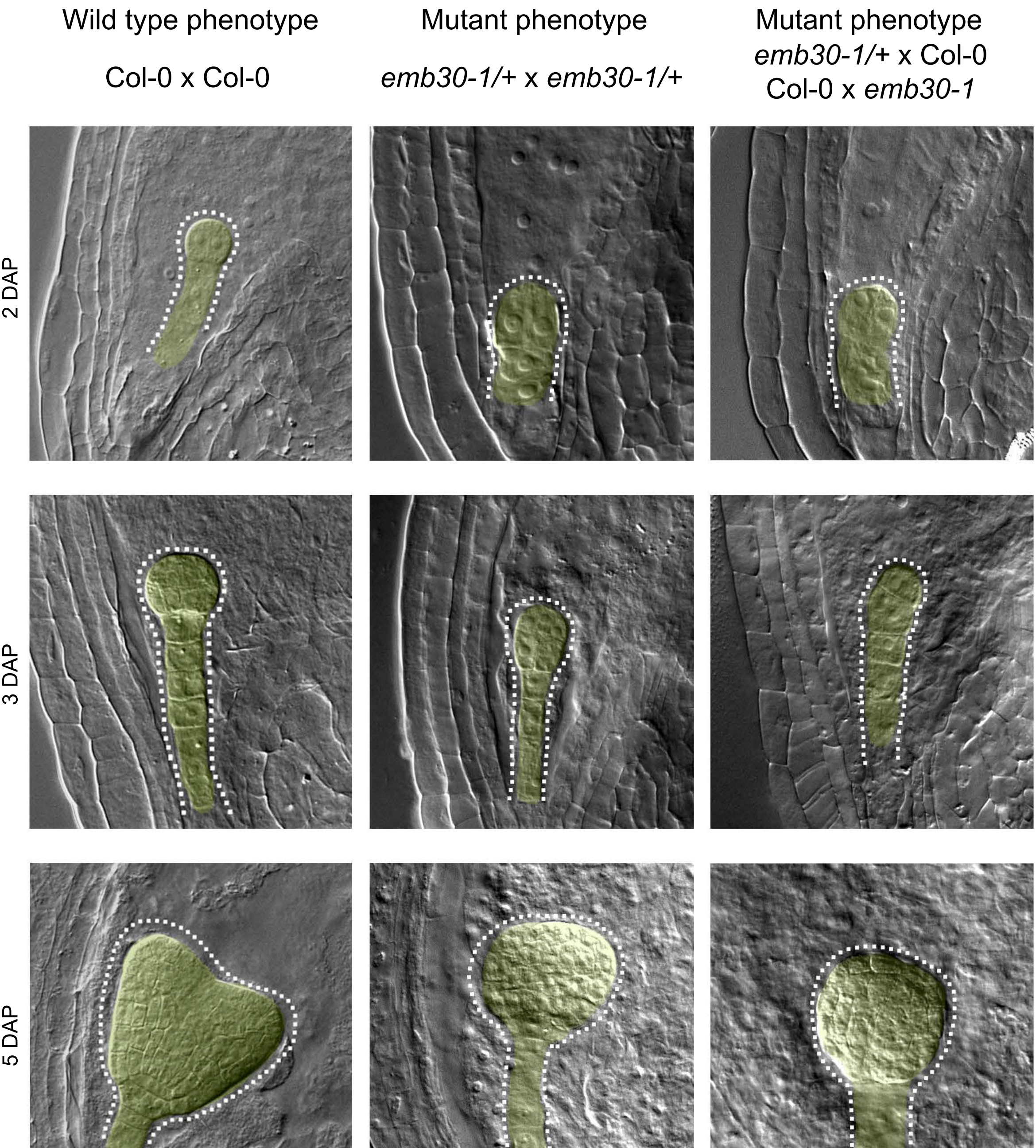


**Protein predicted function:** Encodes a GDP/GTP exchange factor for small G-proteins of the ADP ribosylation factor (RAF) class, and as regulator of intracellular trafficking.

**Seed stock number:** CS6320

### *IYO*

## At4g38440

| Wild type phenotype | Mutant phenotypes | Mutant phenotypes |
|---|---|---|
| Col-0 x Col-0 | *iyo-2/+* x *iyo-2/+* | *iyo-2/+* x Col-0<br>Col-0 x *iyo-2/+* |

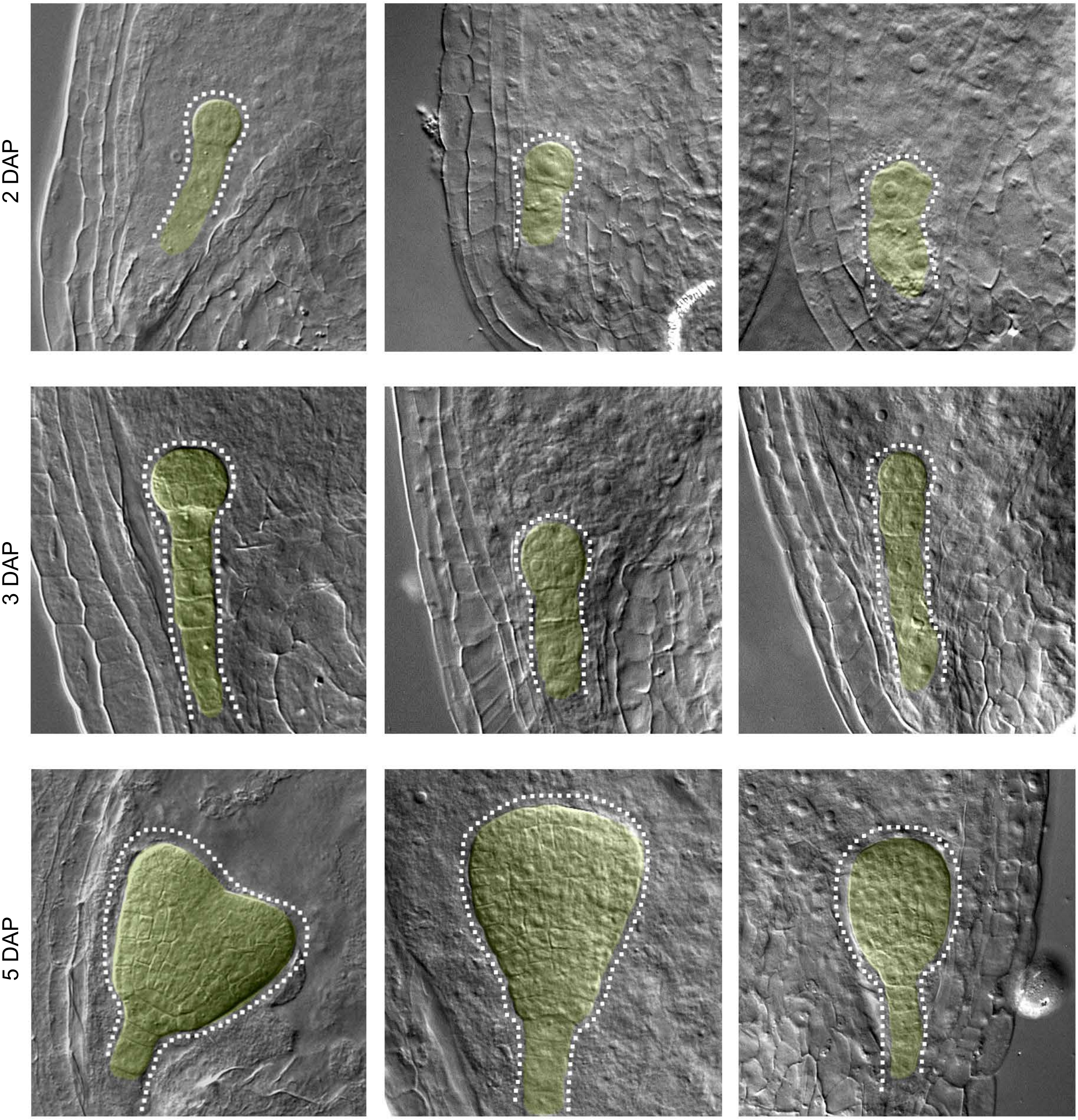


**Protein predicted function:** Encodes MINIYO, a positive regulator of transcriptional elongation that is essential for cells to initiate differentiation.

**Seed stock number:** SALK_099872

### *MIRO1 / EMB2473*

## At5g27540

| Wild type phenotypes | Mutant phenotypes | Mutant phenotypes |
|---|---|---|
| Col-0 x Col-0 | *miro1-1/+* x *miro1-1/+* | *miro1-1/+* x Col-0<br>Col-0 x *miro1/+* |

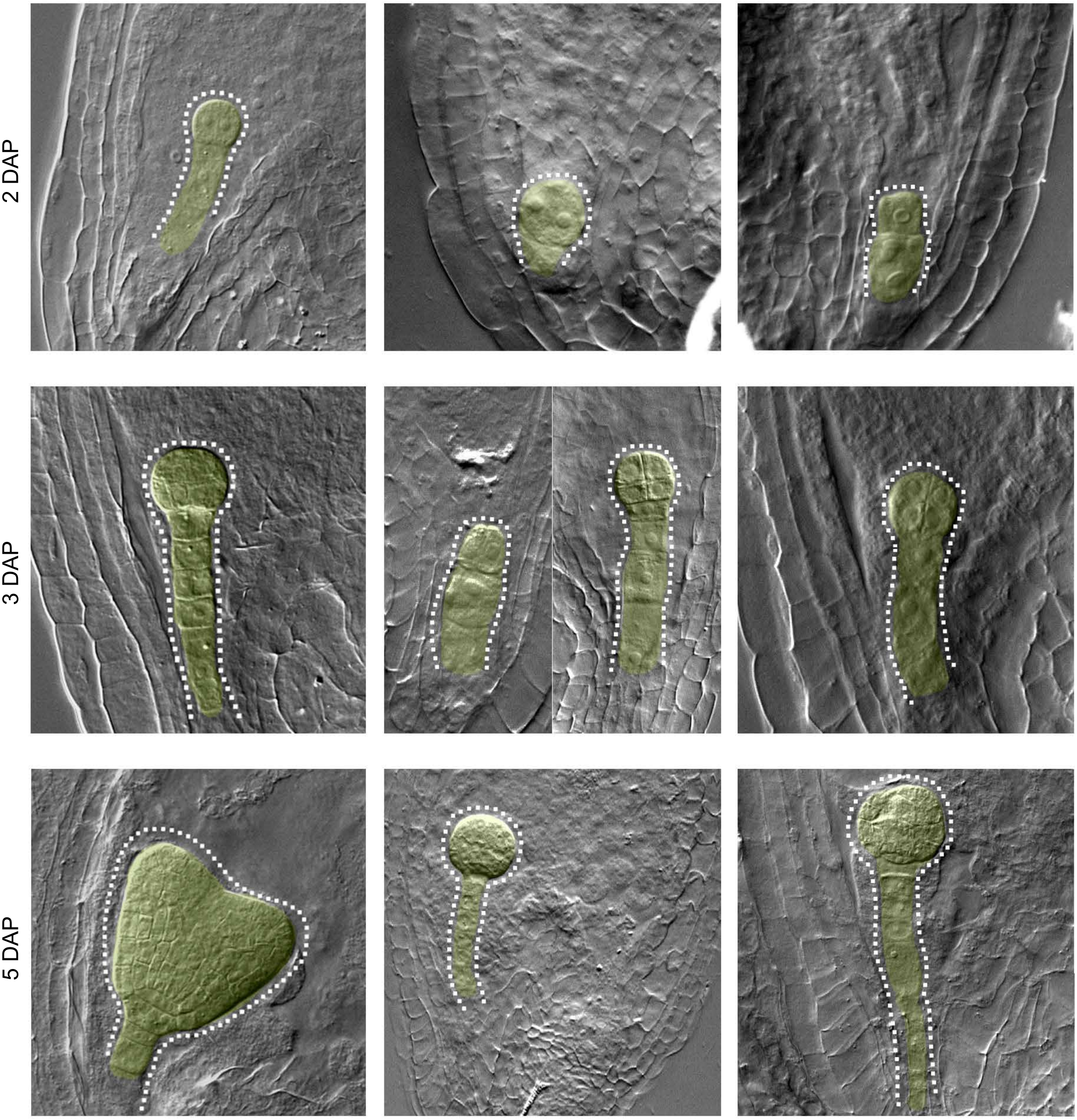


**Protein predicted function:** Encodes a protein with similarity to GTPases that is localized to the mitochondrion.

**Seed stock number:** CS16182

### *MP/ ARF5*

## At1g19850

| Wild type phenotypes | Mutant phenotypes | Mutant phenotypes |
|---|---|---|
| Col-0 x Col-0 | *mp-12/+* x *mp-12/+* | *mp-12/+* x Col-0<br>Col-0 x *mp-12/+* |

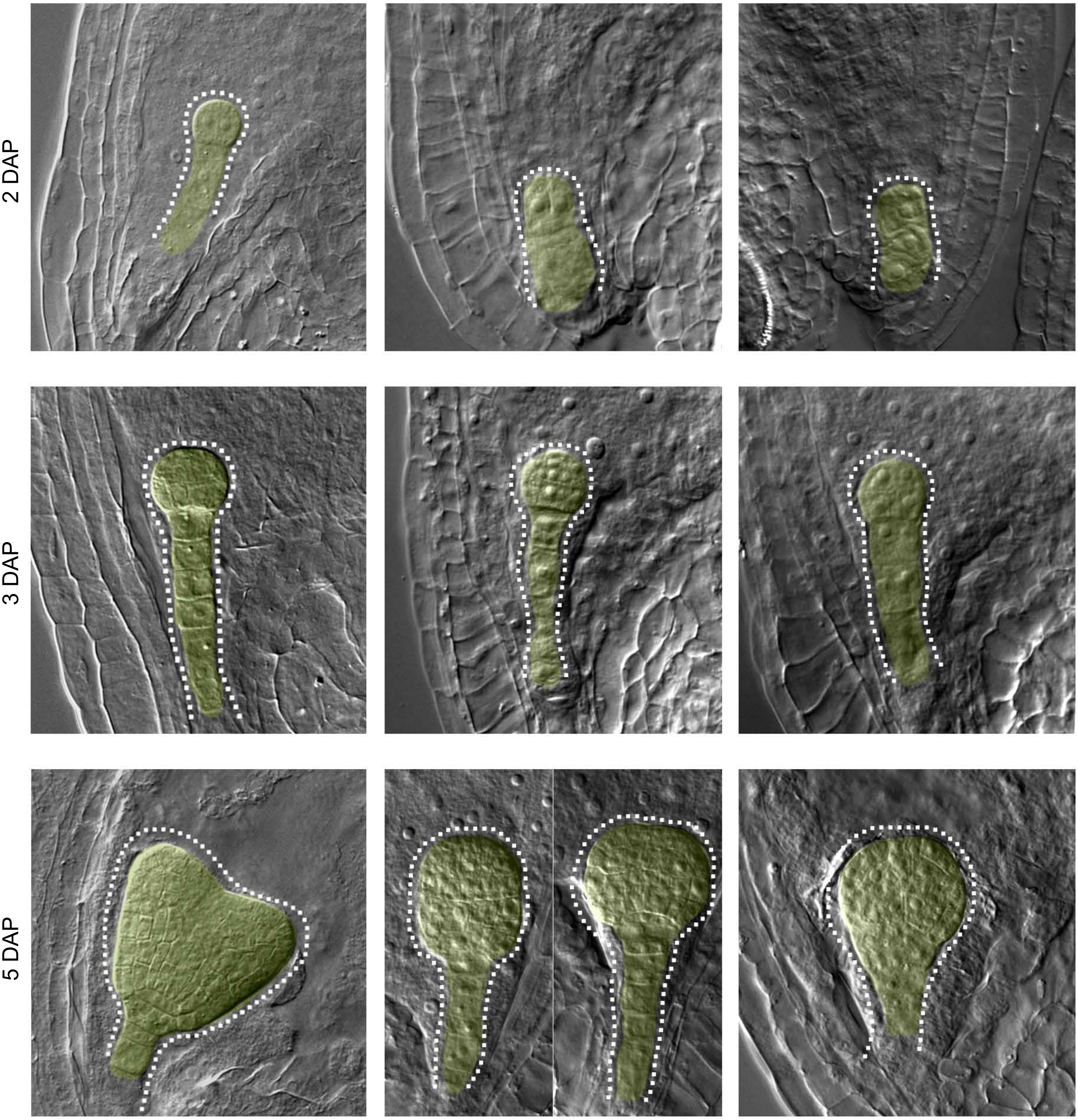


**Protein predicted function:** Encodes a transcription factor (ARF5) mediating embryo axis formation and vascular development.

**Seed stock number:** SALK_149553

### *NAGLU / CYL1*

## At5g13690

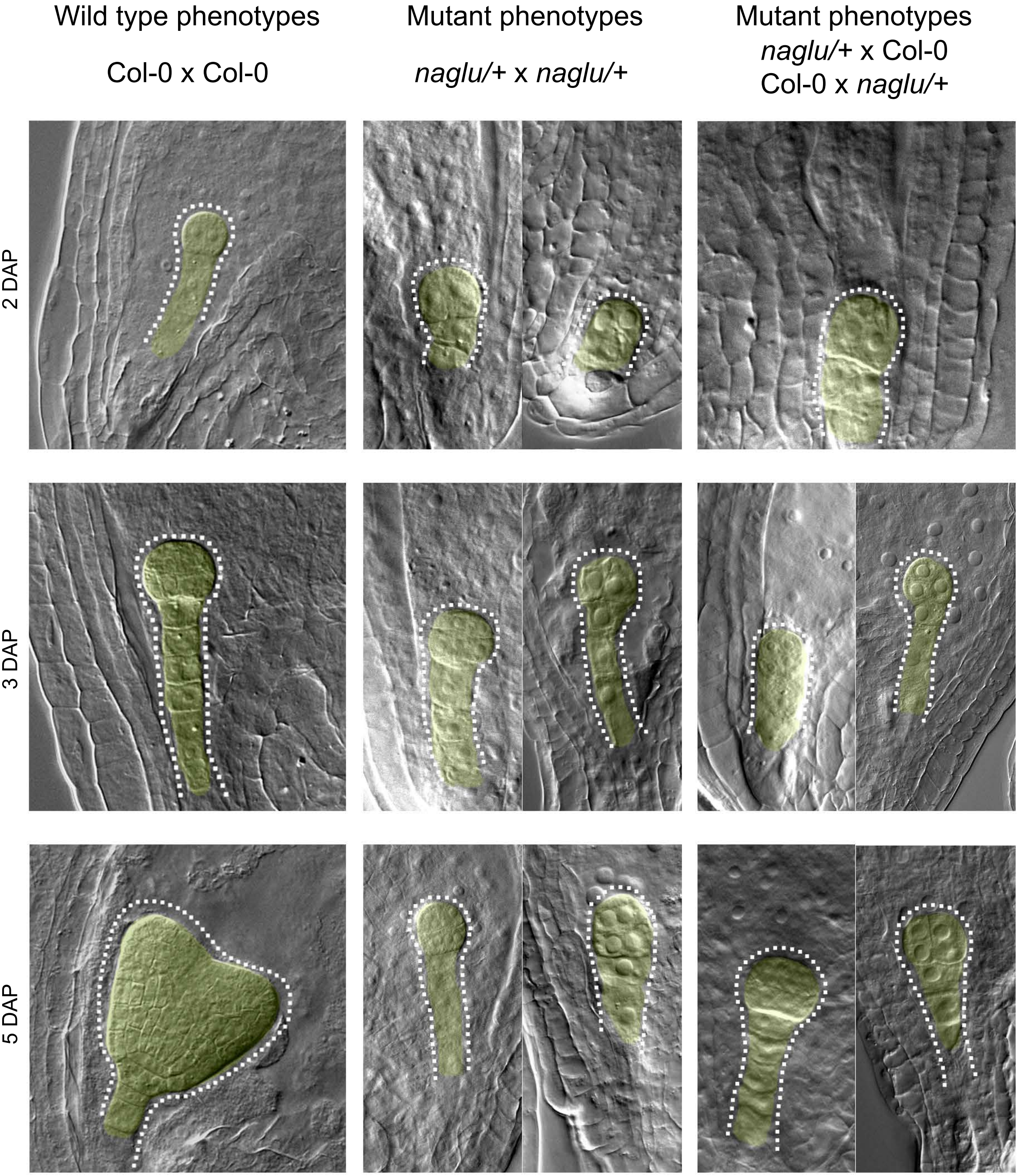


**Protein predicted function:** Encodes an enzyme that is predicted to act as an alpha-N-acetylglucosaminidase (NAGLU).

**Seed stock number:** SALK_065889C

### *NSE1 / EMB1379*

## At5g21140

| Wild type phenotypes | Mutant phenotypes | Mutant phenotypes |
|---|---|---|
| Col-0 x Col-0 | *nse1-2/*+ x *nse1-2/*+ | *nse1-2/*+ x Col-0<br>Col-0 x *nse1-2/*+ |

2 DAP

3 DAP

5 DAP

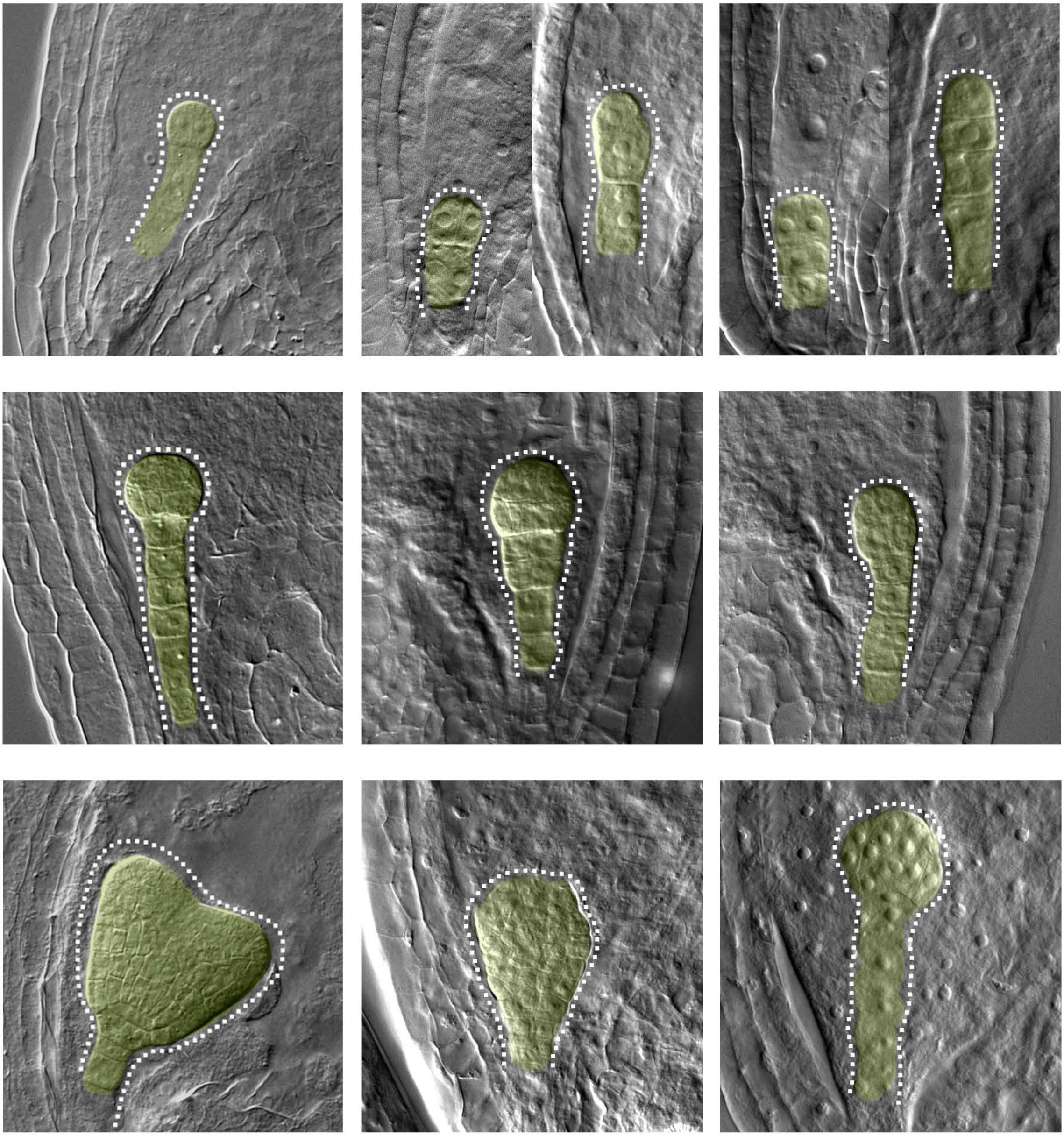

**Protein predicted function:** Encodes a nuclear localized, structural subunit of the SMC 5/6 complex and a non- SMC element.

**Seed stock number:** CS16151

### *NSE3*

## At1g34770

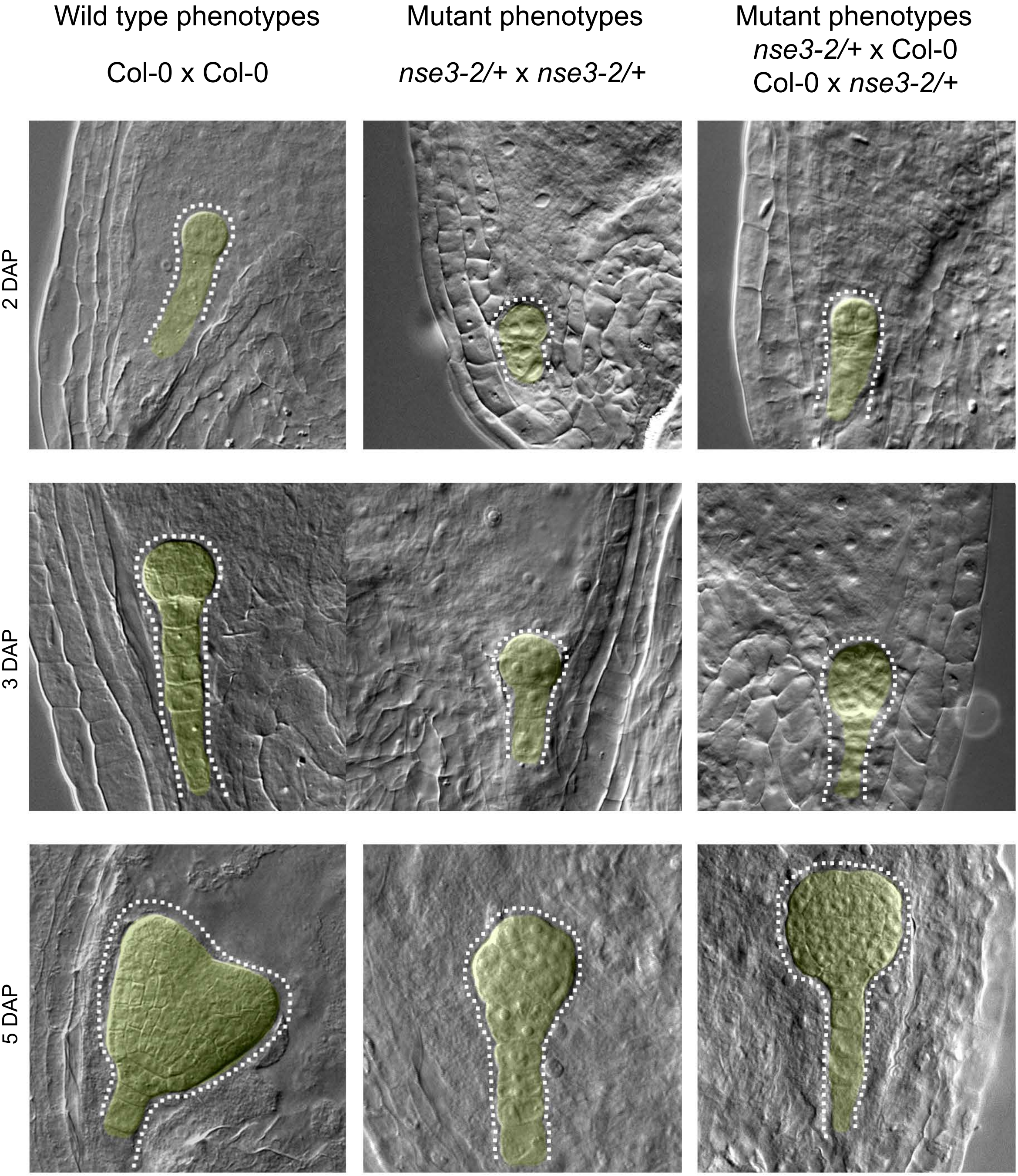


**Protein predicted function:** Encodes a nuclear localized, structural subunit of the SMC 5/6 complex and a non- SMC element.

**Seed stock number:** CS334183

### *PECT1*

## At2g38670

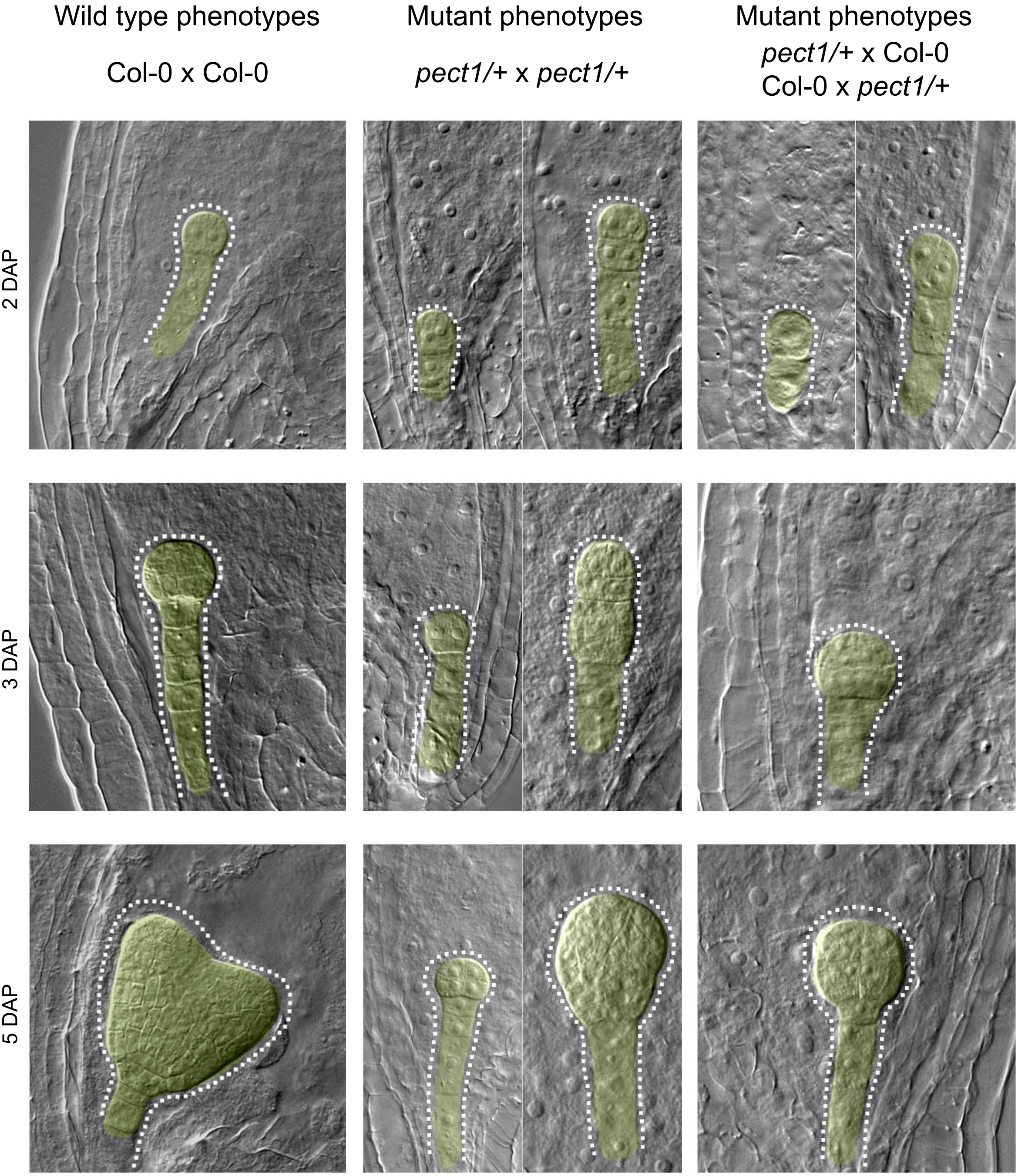


**Protein predicted function:** Encodes a mitochondrial ethanolamine-phosphate cytidylyltransferase, involved in phosphatidylethanolamine (PE) biosynthesis.

**Seed stock number:** SALK_085102

### *PLC2*

## At3g08510

Wild type phenotypes

Col-0 x Col-0

Mutant phenotypes

*plc2-2/+* x *plc2-2/+*

Mutant phenotypes

*plc2-2/+* x Col-0

Col-0 x *plc2-2/+*

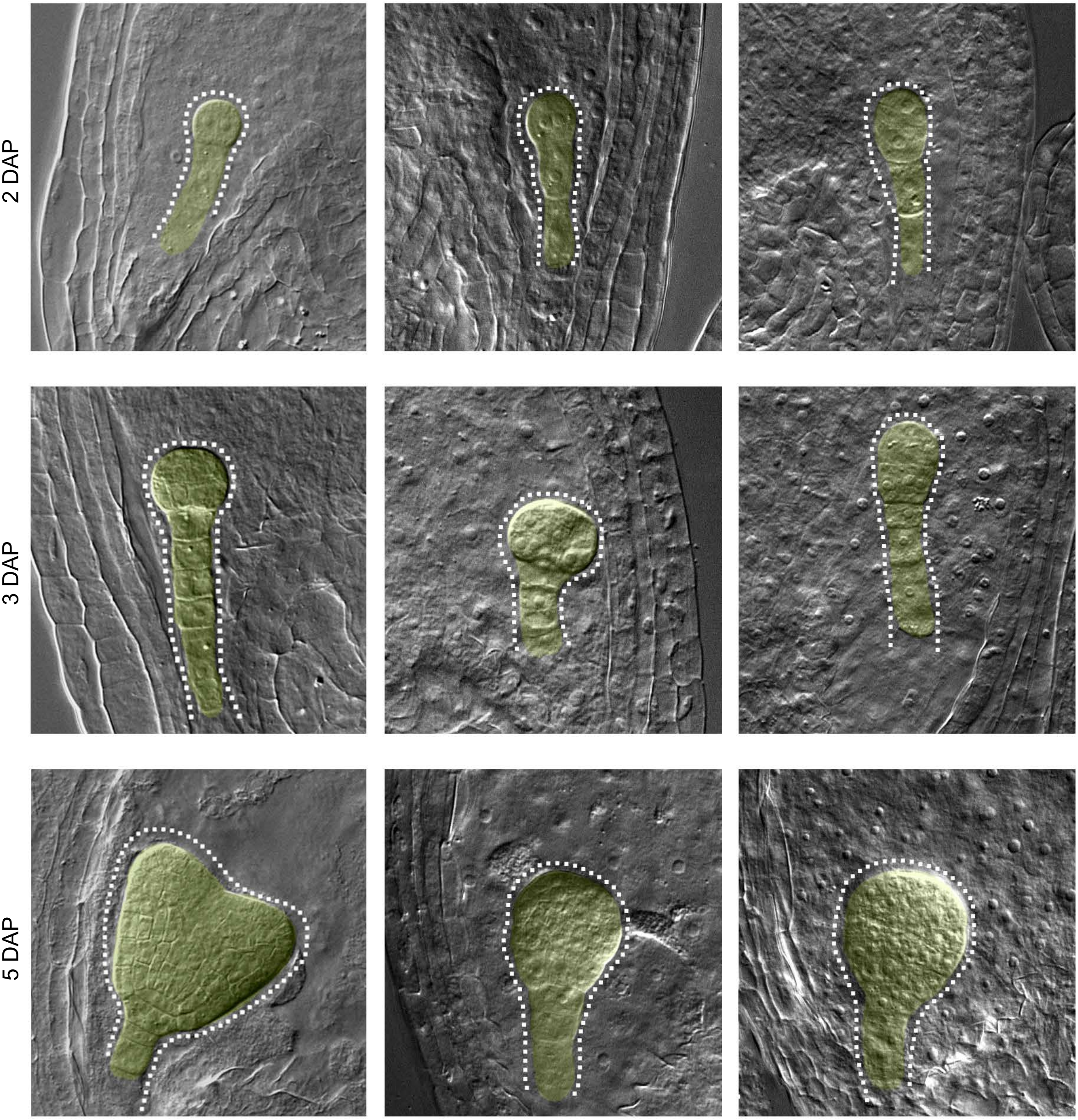


**Protein predicted function:** Phosphoinositide-specific phospholipase C (PI-PLC), catalyzes hydrolysis of phosphatidylinositol 4,5-bisphosphate into inositol 1,4,5-trisphosphate and diacylglycerol.

**Seed stock number:** SALK_152284

### *QQT1 / EMB1705*

## At5g22370

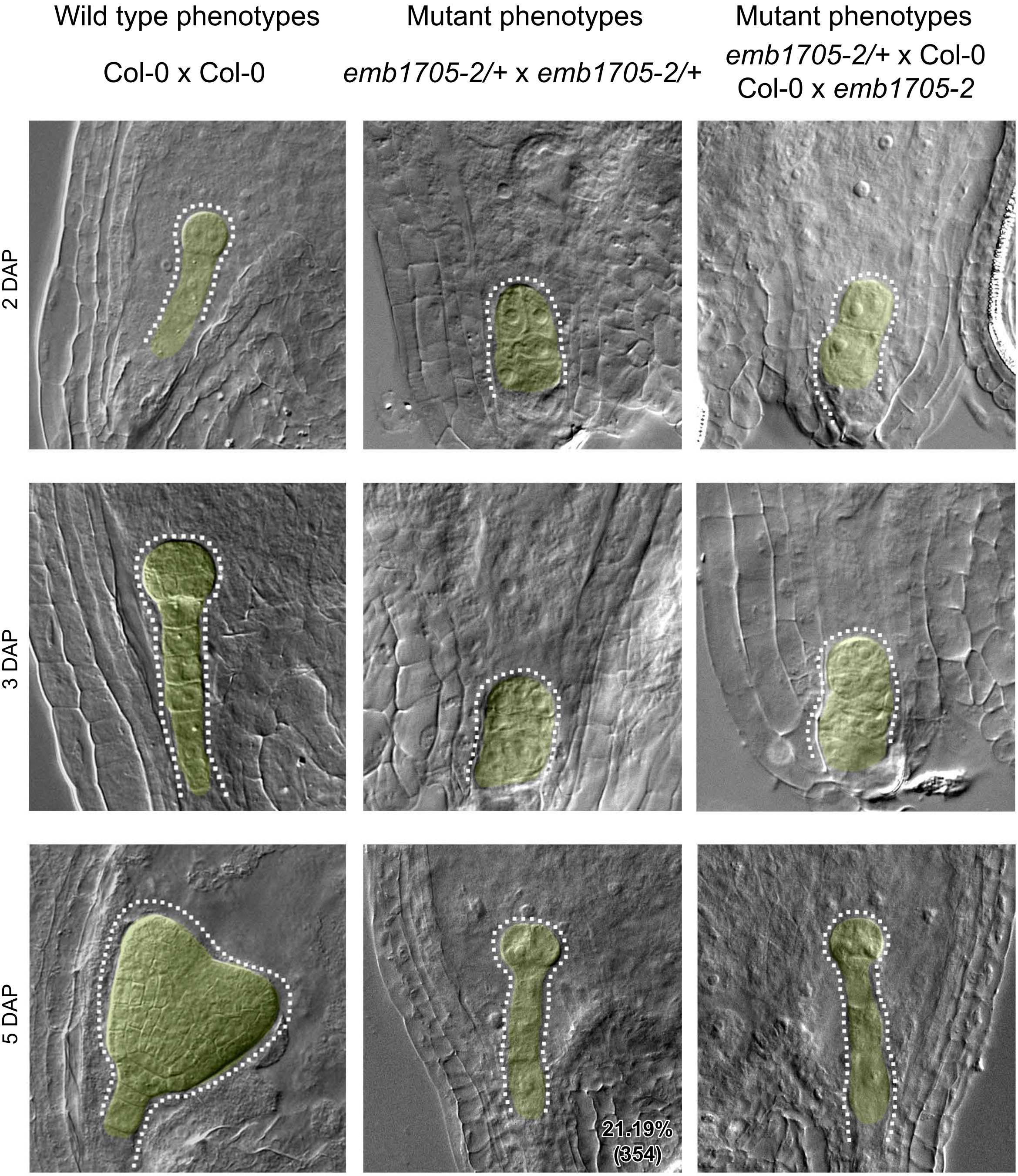


**Protein predicted function:** Microtubule organization.

**Seed stock number:** SALK_059852

### *QQT2*

## At4g21800

| Wild type phenotypes | Mutant phenotypes | Mutant phenotypes |
|---|---|---|
| Col-0 x Col-0 | *qqt2/*+ x *qqt2/*+ | *qqt2/*+ x Col-0<br>Col-0 x *qqt2/*+ |

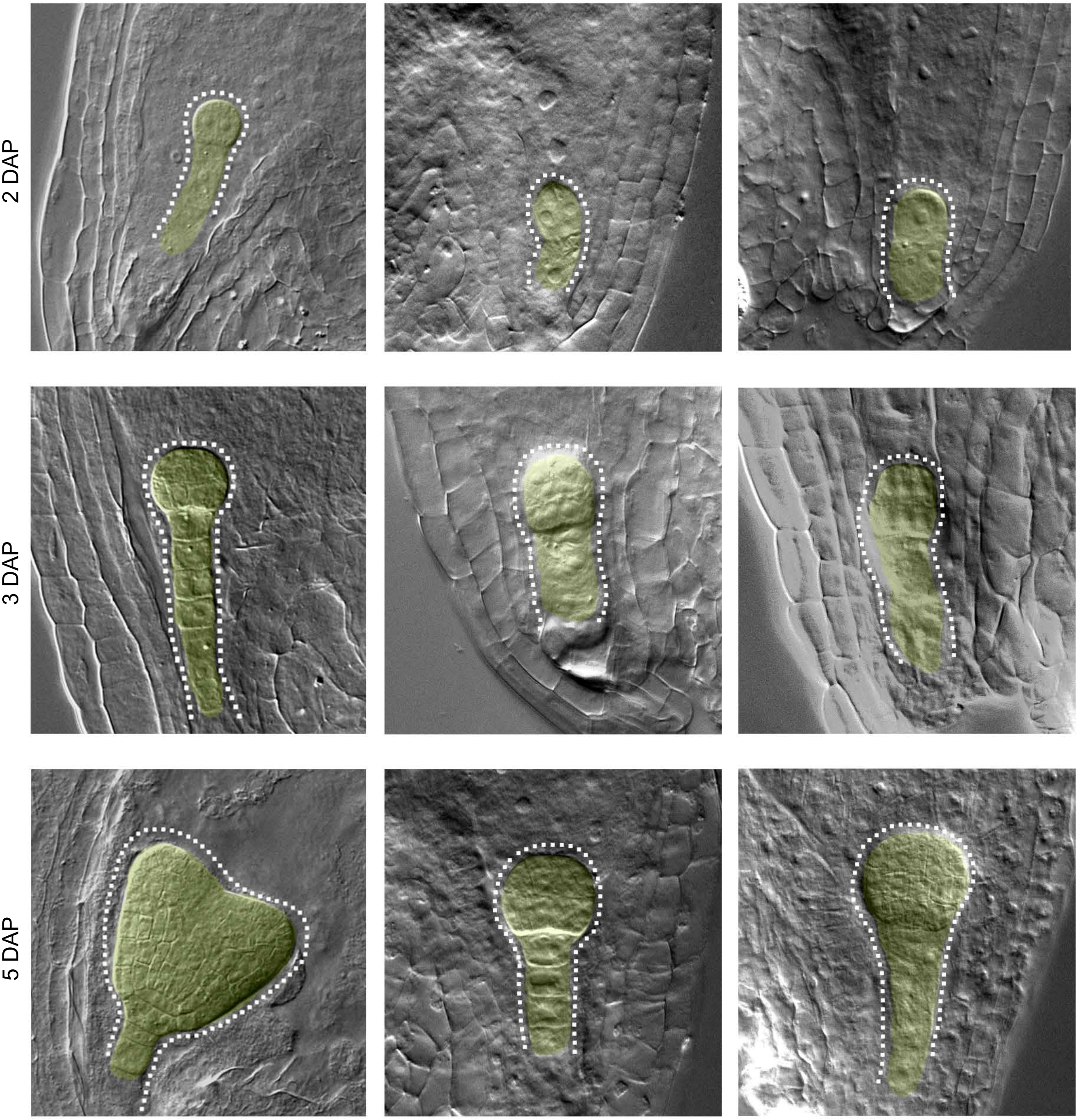


**Protein predicted function:** Microtubule organization.

**Seed stock number:** SALK_026078

### *RFC4 / EMB1968*

## At1g21690

| Wild type phenotypes | Mutant phenotypes | Mutant phenotypes |
|---|---|---|
| Col-0 x Col-0 | *rfc4-1/+* x *rfc4-1/+* | *rfc4-1/+* x Col-0<br>Col-0 x *rfc4-1* |

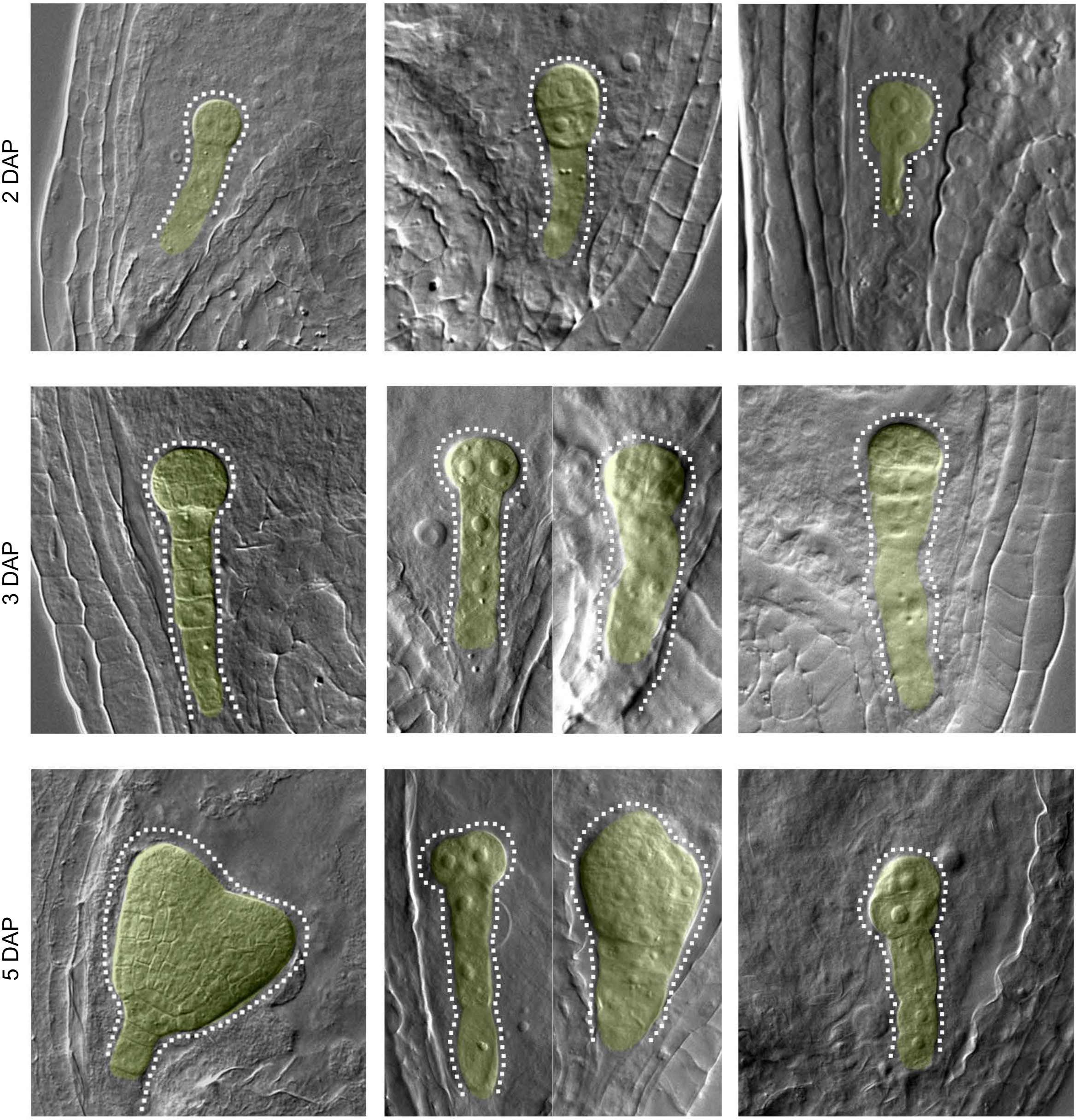


**Protein predicted function:** ATPase family associated with various cellular activities.

**Seed stock number:** CS16161

### *WDR55*

## At2g34260

| Wild type phenotypes | Mutant phenotypes | Mutant phenotypes |
|---|---|---|
| Col-0 x Col-0 | *wdr55-2/+* x *wdr55-2/+* | *wdr55-2/+* x Col-0<br>Col-0 x *wdr55-2* |

2 DAP

3 DAP

5 DAP

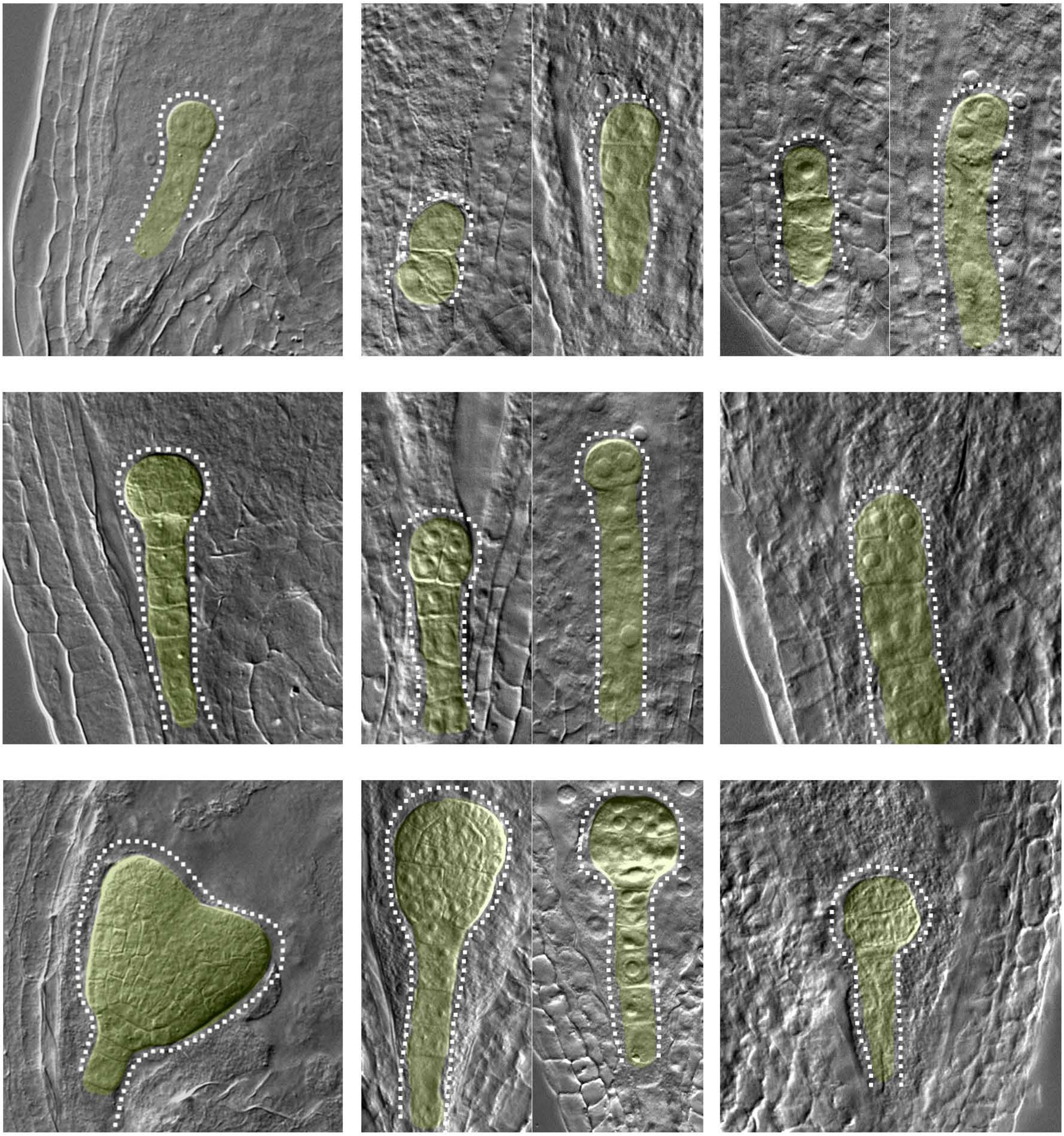

**Protein predicted function:** Encodes a WDxR motif-containing protein that is required for gametogenesis, seed and endosperm development.

**Seed stock number:** CS855491

### *WOX2*

## At5g59340

| Wild type phenotypes | Mutant phenotypes | Mutant phenotypes |
|---|---|---|
| Col-0 x Col-0 | *wox2-1/-* x *wox2-1/-* | *wox2-1/-* x Col-0<br>Col-0 x *wox2-1* |

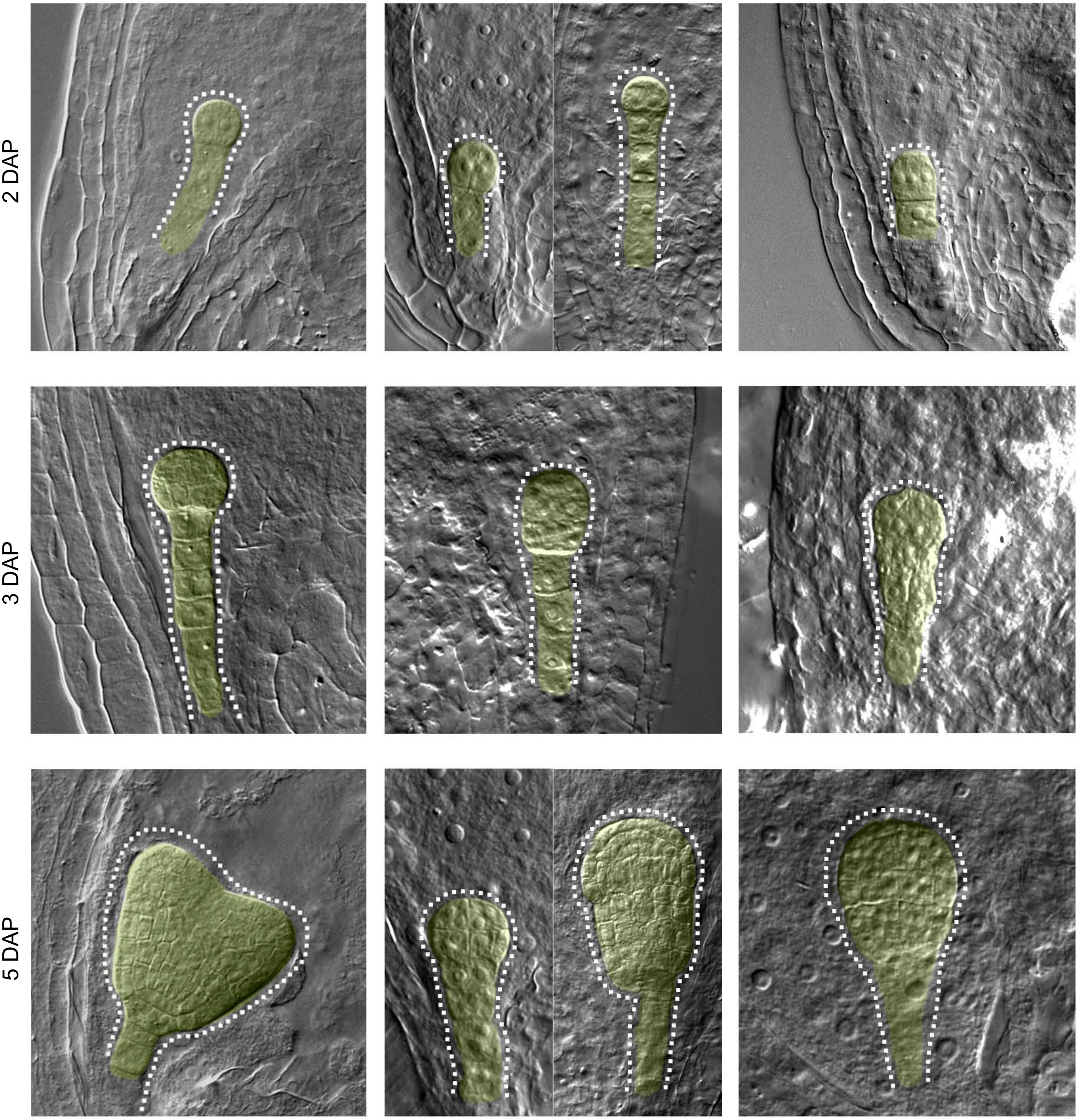


**Protein predicted function:** Encodes a WUSCHEL-related homeobox gene family member with 65 amino acids in its homeodomain.

### *YDA* (Col)

## At1g63700

| Wild type phenotypes | Mutant phenotypes | Mutant phenotypes |
|---|---|---|
| Col-0 x Col-0 | *emb71/+* x *emb71/+* | *emb71/+* x Col-0<br>Col-0 x *emb71/+* |

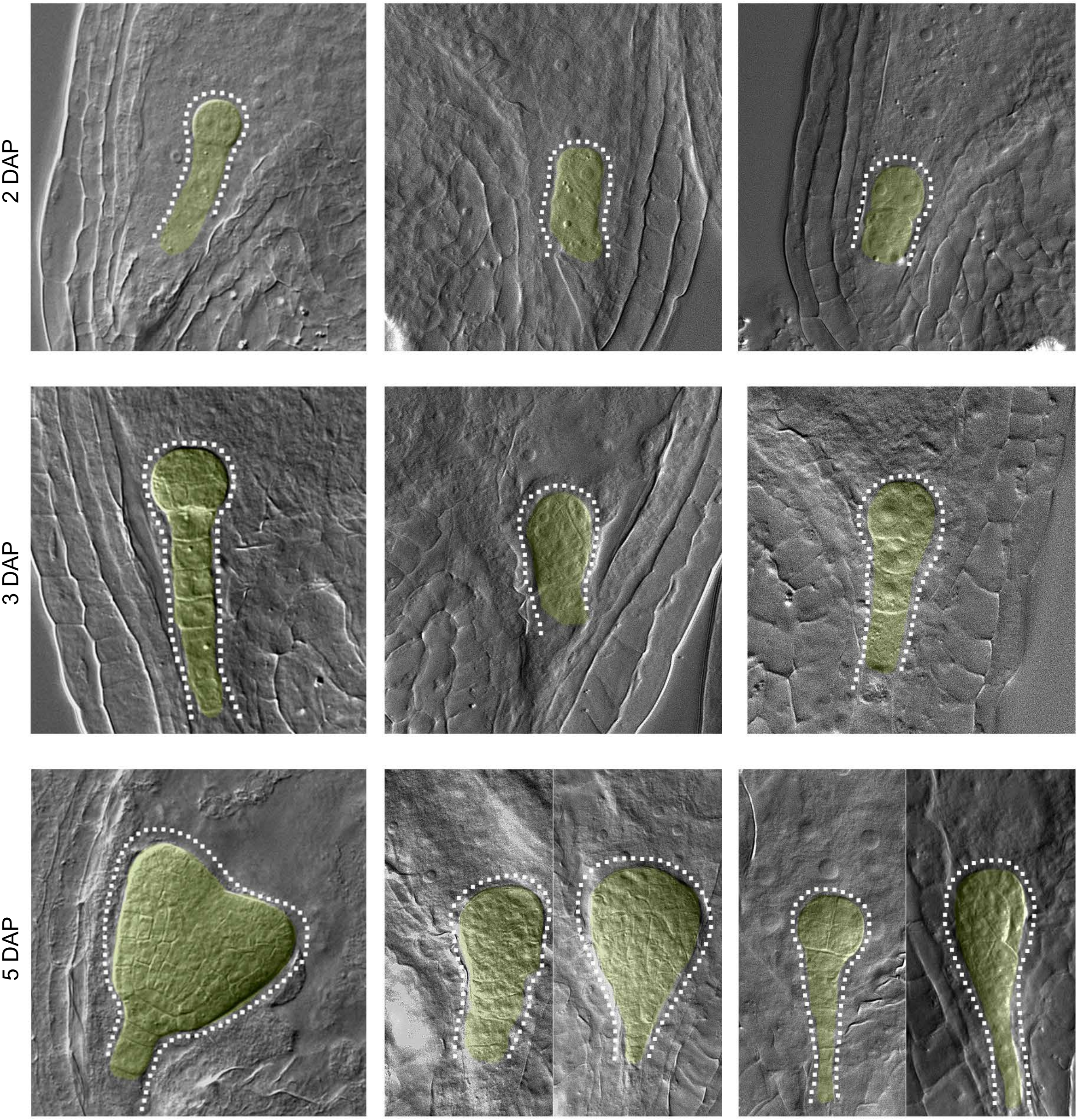


**Protein predicted function:** MAP3K Protein Kinase; Phosphorylation, involved in signal transduction pathways.

**Seed stock number:** CS84618

### *ZYG3*

## At1g07910

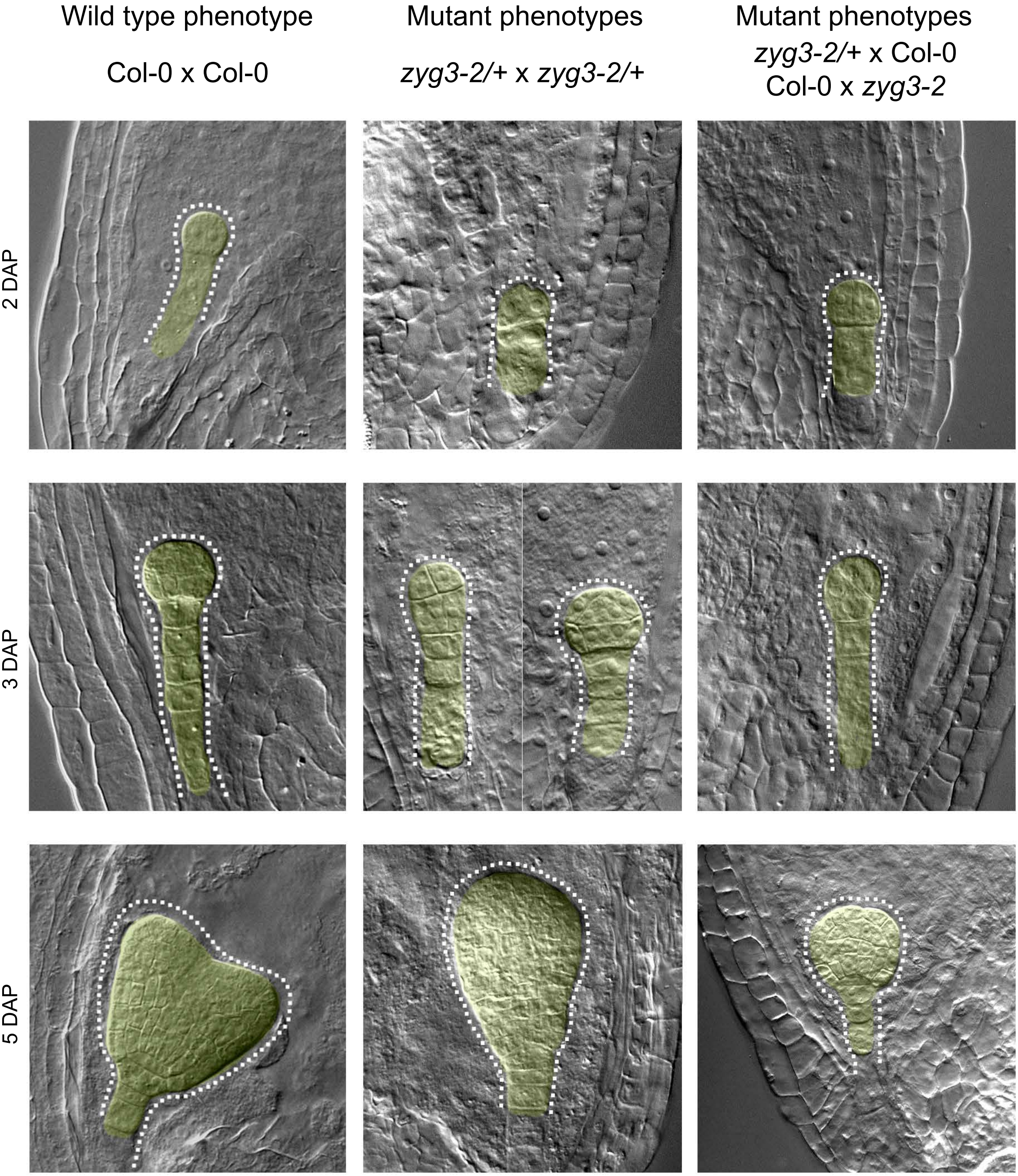


**Protein predicted function:** Encodes a tRNA ligase that resembles the yeast Trl1 RNA ligase in structure and function but very different in sequence.

**Seed stock number:** CS746016
